# Identification and characterization of the surface layer protein AvsA in outer membrane vesicles, antibiotic resistance, and *in vivo* host colonization in *Aeromonas veronii*

**DOI:** 10.1101/2025.04.15.648961

**Authors:** Caroline Vieira Da Silva, Danielle Arsenault, Rebecca Kramer-Earley, Victoria L. Robinson, Kathryn Milligan-McClellan, Joerg Graf

## Abstract

Outer membrane vesicles (OMVs) are important in bacterial communication and the transfer of virulence factors. In this study, we identified and characterized the surface layer protein (SLP) AvsA (*<u>A</u>eromonas <u>v</u>eronii* <u>s</u>urface protein <u>A</u>) in the OMVs of *Aeromonas veronii* Hm21, a strain isolated from the medicinal leech *Hirudo verbana*. The surface layer proteins (SLPs) play critical roles in how bacteria interact with each other and their environments, particularly in mediating antibiotic resistance and facilitating host colonization. Furthermore, we investigate the ability of AvsA to confer protection against antibiotics, affect biofilm formation, and contribute to host colonization, providing insight into antibiotic resistance and two crucial factors contributing to the persistence of the bacteria in its host. Our findings suggest that AvsA enhances antibiotic tolerance, facilitates biofilm development, and is important for successful colonization of the leech digestive tract. These data demonstrate that AvsA performs important roles in a wide range of critical phenotypes. This work provides insights into the functional significance of SLPs in *A. veronii* and highlights AvsA as a potential target for modulating bacterial colonization and resilience against antibiotics.

**Importance:** Outer membrane vesicles (OMVs) are important for bacterial communication, pathogenesis, and stress adaptation, yet how this is accomplished remains poorly understood. Here, we identify and characterize a surface layer protein (SLP), AvsA, associated with OMVs in *Aeromonas veronii* Hm21. Bioinformatic, phylogenetic, and mass spectrometry analyses suggest that *Aeromonas veronii* ORF M001_06550 encodes a surface layer protein (SLP) with high similarity to a characterized *A. hydrophila* SLP, supporting its designation as *Aeromonas veronii* Surface Protein A (AvsA). AvsA forms a paracrystalline layer on bacterial cells and OMVs. Functionally, AvsA contributes to antibiotic resistance, enhances biofilm formation, and is essential for colonization in a symbiotic host, the medicinal leech. These findings highlight a novel role of SLPs in bacterial physiology and host interactions. Given the widespread presence of AvsA homologs, our study provides insights into conserved bacterial mechanisms that may be relevant for both pathogenic and beneficial host-microbe interactions.

## Introduction

Outer membrane vesicles (OMVs) are bilayered structures naturally released by bacteria, playing essential roles in bacterial survival, pathogenesis, and environmental interactions.

Initially identified in Gram-negative bacteria, OMVs are derived from the budding of the outer membrane and serve as vehicles for the transport of biologically active cargo, including lipids, proteins, nucleic acids, and small molecules [1], [2], [3], [4], [5]. The molecular cargo carried by these vesicles reflects their versatile functions, from facilitating horizontal gene transfer (HGT) and interbacterial competition to immune modulation and virulence factor delivery [6], [7] [8], [9]. OMVs have also been implicated in the protection of antibiotic resistance factors, including β-lactamases, efflux pump components, and resistance-conferring genes, and contributing to the rapid dissemination of resistance traits across bacterial populations [10], [11], [12], [13]. Here, we identified an external layer associated with a bacterial OMV and later discovered it to be a surface layer protein (SLP), which was maintained throughout the vesiculation process.

Surface layer proteins (SLPs) are highly organized crystalline arrays of protein or glycoprotein subunits found on the exterior of many bacterial and archaeal cells [14], [15], [16], [17]. SLPs function as a protective barrier, mediate host interactions, and contribute to environmental adaptation [17]. Their structural composition and arrangement can vary significantly between organisms, suggesting an adaptation to specific ecological niches or host environments [18]. The presence of SLPs is widespread among pathogenic, environmental, and symbiotic bacteria, enhancing bacterial fitness in various niches. In pathogenic and environmental bacteria, SLPs contribute to immune evasion by masking immunogenic components or inhibiting host immune responses [19]. Additionally, they facilitate adhesion to host tissues and biofilm formation, enabling resilience in harsh conditions and aiding nutrient acquisition [20]. Similarly, in symbiotic bacteria, SLPs are crucial for establishing and maintaining associations with their hosts, often mediating attachment or providing protection from host immune factors [21], [22].

The genus *Aeromonas* encompasses a group of Gram-negative, rod-shaped bacteria widely distributed in aquatic environments. This genus includes several pathogenic species, which are capable of infecting both poikilothermic and homoeothermic hosts, causing a range of infections such as gastroenteritis, wound infections, and septicemia [23], [24]. The SLP in *Aeromonas hydrophila* has been associated with its ability to evade host immune defenses and establish infections, highlighting its role in virulence [25], [26]. *Aeromonas veronii* is particularly notable for its pathogenicity in both humans and animals [27], [28] and beneficial associations with fish and leeches [29], [30]. It is frequently isolated from cases of septicemia in fish and gastroenteritis in humans [27], is shown to influence the development of zebrafish digestive tracts, and releases hemolysins in blood-feeding leeches [31], [32], [33]. Here we describe the SLP of *A. veronii* Hm21, a strain isolated from the medicinal leech *Hirudo verbana* [34]. This strain has evolved to thrive in the leech gut, where it plays a symbiotic role, assisting in the digestion of blood meals and helping the leech resist colonization by other pathogens [34], [35].

While SLPs have been described in a few *Aeromonas* strains, their role in *Aeromonas veronii* and in symbiotic interactions has not been reported. *Aeromonas* SLPs can act as adhesins, facilitating bacterial adhesion to host tissues and surfaces, a critical first step in infection [36], [37]. Additionally in other bacteria, these proteins may interact with host immune cells to evade detection or modulate immune responses, further enhancing bacterial pathogenicity [2], [38], [39]. We hypothesize that in *A. veronii* Hm21, the surface layer protein, AvsA (*<u>A</u>. <u>v</u>eronii* <u>s</u>urface layer protein <u>A</u>), is essential for both bacterial defense and host interactions. In this study, we investigate the role of AvsA in mediating resistance to the antibiotics tobramycin and ciprofloxacin, which are commonly used to prevent or treat *Aeromonas* infections after leech therapy. Additionally, we examine the importance of AvsA in host colonization, assessing its function in enabling *A. veronii* to proliferate in the leech gut environment and in biofilm formation, which is essential for establishing symbiotic and pathogenic interactions [21], [40].

## Results

### Characterization of *A. veronii* outer membrane vesicles

Outer membrane vesicles (OMVs) were isolated from HM21RS, a spontaneous rifampicin (R) and streptomycin (S)-resistant derivative of *A. veronii* Hm21. OMVs were isolated through ultracentrifugation followed by an Optiprep gradient purification method [41], [42]. To confirm the isolation of OMVs, a 12% sodium dodecyl sulfate-polyacrylamide gel electrophoresis (SDS-PAGE) was performed, revealing two prominent bands at approximately 47 kDa and 30 kDa, along with a less prominent band around 37 kDa, suggesting the presence of specific proteins that are highly abundant in the vesicles (Fig. 1A). Further structural analysis of the OMVs using transmission electron microscopy (TEM) revealed an unusual vesicle morphology with a defined external surface layer covering the outer membrane (Fig. 1B). To further characterize the protein composition of the OMVs, proteomics analysis using liquid chromatography-mass spectrometry (LC-MS/MS) was performed on OMV samples. This analysis identified 41 proteins, including uncharacterized proteins, flagellins, elongation factors, ribosomal proteins, and other functional and structural components commonly associated with bacterial OMVs [43], [44] (Sup. Table 1). The protein with the highest spectral count indicating its abundance in the OMVs was an uncharacterized protein (M001_006550) with a molecular weight of 47 kDa (Table 1).

**Figure 1.**
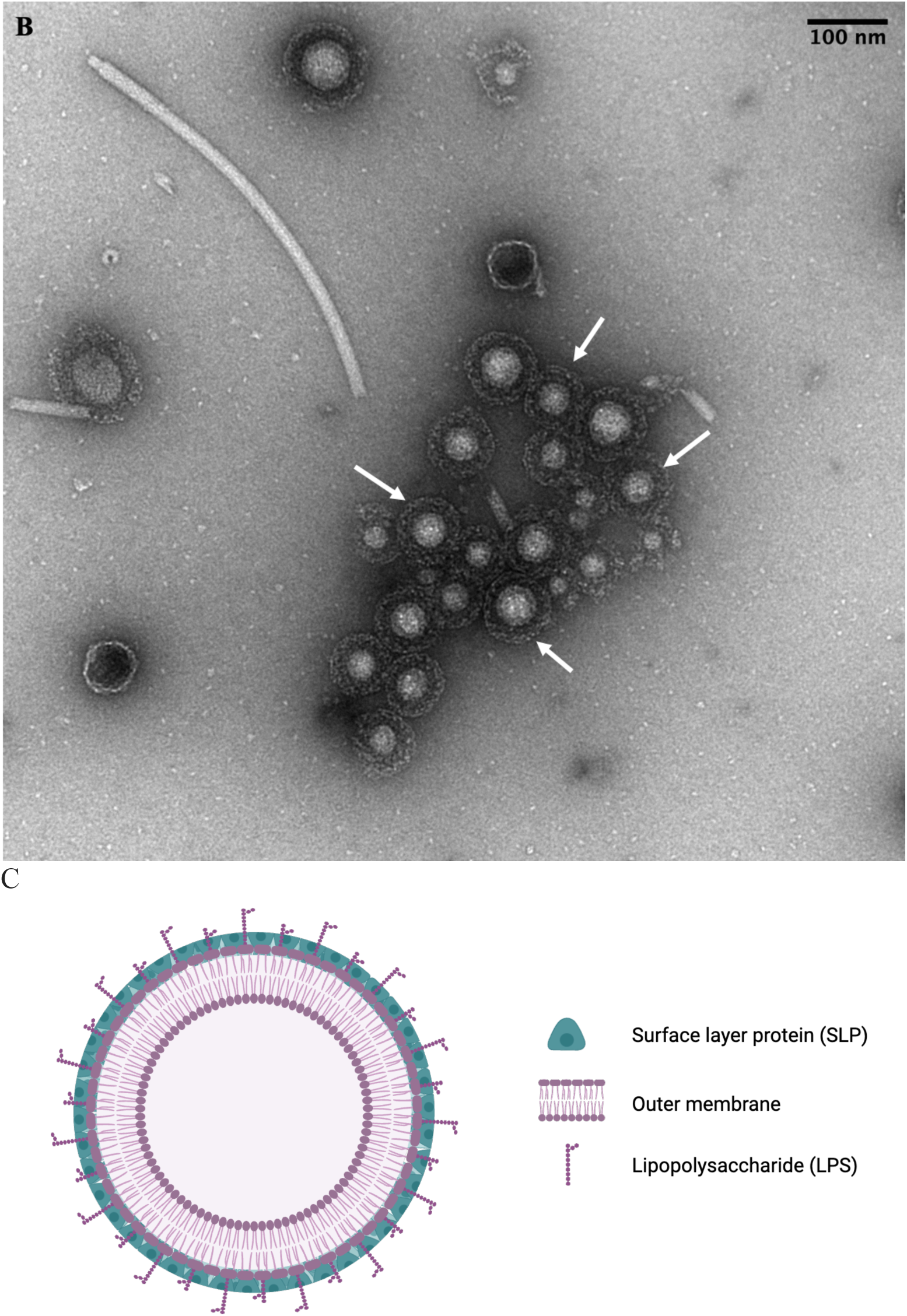
Detection of surface layer protein on outer membrane vesicles of *A. veronii* Hm21. **(A)**. A very abundant protein was detected in the wild-type OMVs. Samples were loaded into a 12% SDS-PAGE. Lanes 1 and 8 contain the protein ladder (sizes indicated in kDa on the left). Lanes 2 through 7 show Hm21RS OMVs fractions of Optiprep (OP) gradient. A prominent band at approximately 47 kDa is observed in the 50% OP (lane 7) in the Hm21RS OMVs sample. Furthermore, a prominent band at ∼30 kDa was observed. **(B)** TEM micrograph shows a crystalline layer (white arrows) around the OMVs isolated from the Hm21RS strain. Scale bar: 100 nm. **(C)** A hypothetical scheme of the OMVs from Hm21 indicates the position of the lipopolysaccharide (LPS) and the AvsA SLP with the outer membrane.

**Table 1.** Top 15 protein composition of OMVs isolated from *A. veronii* Hm21 determined by liquid chromatography-mass spectrometry (LC-MS) and compared to the predicted proteins of Hm21.

| Identified Proteins | Accession<br>Number | Alternate ID | Molecular<br>Weight<br>(kDa) | Total<br>Spectrum<br>Count |
| --- | --- | --- | --- | --- |
| Uncharacterized<br>protein | A0A7Z3TVW6 | M001_006550 | 47 | 272 |
| Lateral flagellin | A0A7Z3YHV7 | LafA | 31 | 28 |
| Lateral flagellin | A0A7Z3YHT0 | LafA | 30 | 16 |
| Elongation factor<br>Tu | A0A7Z3TT05 | Tuf | 43 | 16 |
| Glyceraldehyde-3-phosphate<br>dehydrogenase | A0A7Z3TSE5 | Gap | 35 | 9 |
| Elongation factor G | A0A7Z3TT82 | FusA | 78 | 8 |
| Arginine deiminase | A0A7Z3TT59 | ArcA | 46 | 7 |
| ATP synthase<br>subunit alpha | A0A7Z3TTL0 | AtpA | 55 | 7 |
| Collagenase | A0A7Z3TSN7 | M001_018910 | 104 | 5 |
| 30S ribosomal<br>protein S1 | A0A7Z3TWQ7 | RpsA | 61 | 5 |
| Uncharacterized<br>protein | A0A7Z3TTU3 | M001_000540 | 33 | 5 |
| 30S ribosomal<br>protein S3 | A0A7Z3YHR6 | RpsC | 26 | 5 |
| Fructose-<br>biphosphate<br>aldolase | A0A7Z3TS79 | FbaA | 39 | 4 |
| 50S ribosomal<br>protein L9 | A0A7Z3YI44 | RplI | 15 | 4 |
| Alkyl<br>hydroperoxide<br>reductase C | A0A7Z3TQP0 | AhpC | 21 | 4 |

### Bioinformatic analysis of M001_006550 reveals similarity to a surface layer protein

Based on the mass spectrometry data, we focused our bioinformatic analysis on open reading frame (ORF) M001_06550. A search against NCBI’s non-redundant protein sequence database using the amino acid sequence of ORF M001_006550 as query retrieved 152 unique sequences with significant similarity (e-value ≤ 0.006), representing 312 genomes in total. Except for one sequence, all hits were annotated as hypothetical proteins. One sequence, AAA67043.1, with an e-value of 0.0, 100% query cover and an identity of 81.66% to M001_006550 was from a characterized paracrystalline surface layer protein (SLP) from *A. hydrophila.* The main differences are clustered between amino acid 80 and 120, including a 10 amino acid deletion in *A. veronii.* The presumptive secretion signal sequence at the N-terminus is 18 aa in *A. veronii* and 21 in *A. hydrophila,* but this difference could be due to selecting a different start codon. This SLP was well characterized by the research group from Trevor Trust [38], [45]. From all significant search matches, sequences longer than 300 amino acids were aligned to construct a maximum-likelihood phylogeny. The sequence from *A. veronii* Hm21 belongs to clade of 18 sequences, which included the *A. hydrophila* SLP, and range in sequence identify from 73% to 99% with sequences originating from *A. allosaccarophila, A. bestiarum, A. caviae, A. dhakensis, A. diversa, A. hydrophila, A. jandaei, A. salmonicida, A. schubertii,* and *A. veronii* (Fig. 2B). The sequences were obtained from the genus *Aeromonas* (230 sequences) and the genera *Stutzerimonas* (33)*, Azotobacter* (13)*, Alteromonas* (4)*, Azorhizophilus* (4)*, Marinobacter* (3), *Pseudomonas* (3)*, Bowmanella* (2), *Paucibacter* (2)*, Aestuariibacter* (1)*, Ectopseudomonas* (1)*, Enterovibrio* (1)*, Rubrivivax* (1), and *Usitatibacter* (1). The genera were ranked according to increasing e-values and additional sequences were from strains not identified to the genus level. It should be noted that all but nine of the unique sequences had an e-value ≤ 1e-10 and all but six sequences had a coverage of ≥ 50%. Based on this analysis and biochemical characterization performed on AAA67043.1, which was demonstrated to be a paracrystalline surface layer protein, we hypothesize that M001_06550 encodes a surface layer protein and name the gene *<u>A</u>eromonas <u>v</u>eronii* <u>S</u>urface Protein <u>A</u>, *avsA*.

**Figure 2.**
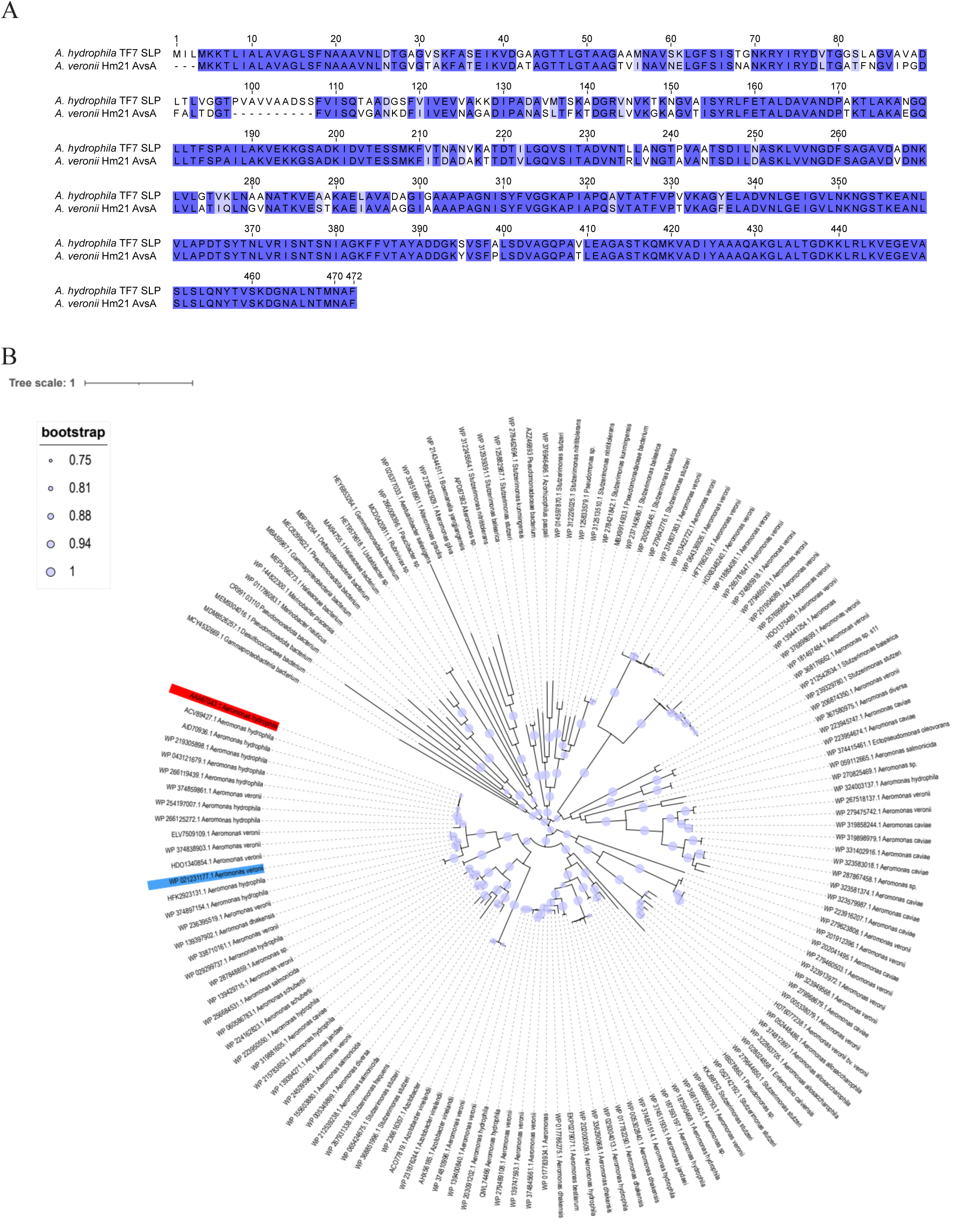
Alignment and Phylogeny of M001_06550 and *A. hydrophila* paracrystalline surface layer protein (SLP). **A.** The sequence of M001_06550 from Hm21 was aligned to the sequence of the well-characterized SLP protein from *A. hydrophila* TF7. The overall identity was 86.77% and the major difference was a 10 amino acid deletion. **B.** A maximum-likelihood phylogeny was calculated from 152 amino acid sequences that had significant sequence similarity (e-value ≤ 0.006 and were at least 300 amino acids long). M001_06550 is shown in blue with the identifier WP 021231177.1. The only annotated protein in this phylogeny is shown in red, AAA67043.1, the paracrystalline SLP of *A. hydrophila.* The bootstrap support is indicated by the size of the circle.

### Deletion of *avsA* affects cell surface and OMVs of *Aeromonas veronii* Hm21RS

In the wild-type strain, *avsA* is 1,380 base pairs (bp) and is situated downstream of the Type 2 Secretion System (T2SS) gene, *pulD*, and upstream of an O-antigen operon gene, *rfbD* (Fig. 3A). Hm21RSΔ*avsA* was created by deleting all but 70 bp of *asvA*, keeping *pulD* and *rfbD* intact (Fig. 3.A).This rendered *avsA* nonfunctional. The complement strain Hm21RSKΔ*avsA*Tn7::*avsA* was constructed by inserting the full-length *avsA*, with the promoter region, into Tn*7*. Successful knockout and complementation were confirmed by PCR.

**Figure 3.**
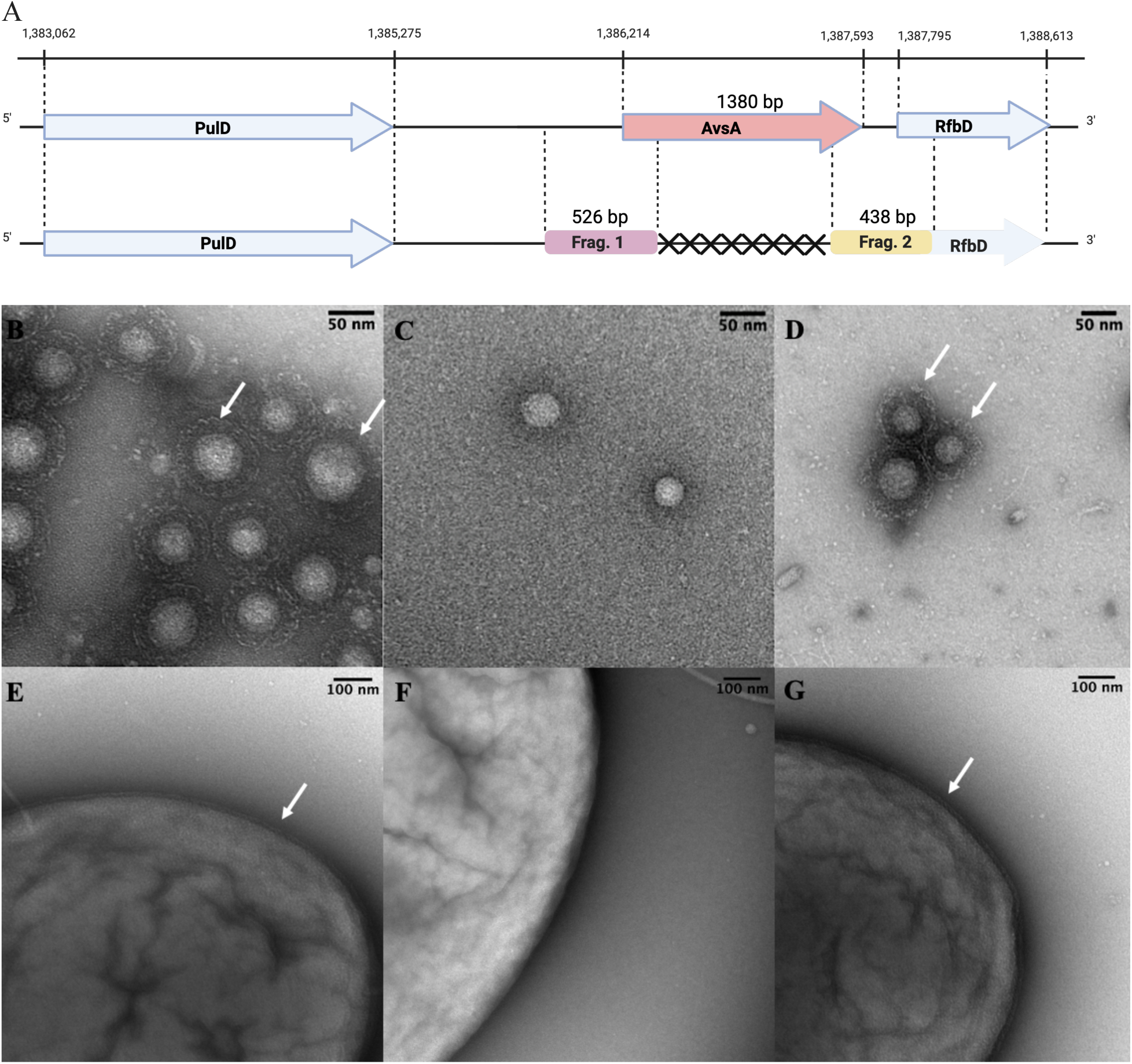
Construction of Hm21Δ*avsA* strain and its impact on cell morphology in *Aeromonas veronii* Hm21. **(A)** Schematic representation of the wild-type (top) and deletion mutant (bottom) genetic context of *avsA*. In the Hm21RS strain, the 1380 bp *avsA* gene is located downstream of the type 2 secretion system (T2SS) *pulD* gene and upstream of the o-antigen protein RfbD. Fragment 1 (526 bp) and Fragment 2 (438 bp) represent the regions targeted for amplification. In the Hm21Δ*avsA* deletion mutant, the *avsA* gene was truncated to 70 bp. **(B-G).** Transmission electron microscopy (TEM) analysis of the bacterial morphology. The presence of the AvsA is indicated with a white arrow. **(B)** A surface layer protein, AvsA, surrounds the OMV membrane. The white arrows indicate the AvsA protein in the parental Hm21RS strain, which exhibits a well-defined external crystalline structure. **(C)** The Hm21RSΔ*avsA* strain, in which the knockout of *avsA* resulted in the loss of the external crystalline structure. **(D)** The complemented strain Hm21RSKΔ*avsA*Tn*7::avsA*, with K indicating resistance to kanamycin, in which the crystalline structure was restored. **(E)** Hm21RS strain shows the AvsA protein (arrow), whereas **(F)** in the Hm21Δ*avsA* strain, the AvsA protein is not observed. **(G)** Complementation with Hm21RSKΔ*avsA*Tn*7::avsA* restores the AvsA protein, resembling the Hm21RS phenotype (arrows). Scale bars: 50 nm.

Transmission electron microscopy (TEM) analysis revealed a distinct difference in the cell surface structures when AvsA was deleted in whole cells and OMVs. While the parent cell had an apparent crystalline outer layer covering its OMVs and outer membrane (Fig. 3B and 3E), the mutant lacked this layer on the OMVs and cells (Fig. 3C and Fig. 3F). In the complement (Hm21RSKΔ*avsA*Tn*7::avsA*), the crystalline surface layer was detected (Fig. 3D and 3G), thus linking the crystalline surface layer to a functional AvsA. Based on these results, we propose that AvsA coats the OMVs with a paracrystalline surface layer (Fig. 3H).

### AvsA protects *A. veronii* Hm21 against Tobramycin and Ciprofloxacin

The surface layer protein has been implicated in conferring protection against environmental stressors, including antimicrobial agents [46], [47]. The contribution of AvsA to the protection against antibiotics that target Gram-negative bacteria, including kanamycin; tetracycline; gentamicin; tobramycin (To); ciprofloxacin (Cp); and polymyxin B and polymyxin E, was tested in strains Hm21RS, Hm21RSΔ*avsA*, and Hm21RSKΔ*avsA*Tn*7::avsA* strains. Traditional growth curves and E-strip test analysis showed no significant phenotypic differences between the strains, so we performed a more sensitive assay, the time-kill assay (Suppl. Table 1, Suppl. Figures 1 and 2), on To and Cp.

For the susceptibility testing, we selected concentrations below and above the minimum inhibitory concentration of the strain tested. For To, we used concentrations of 0.5, 5, 10 and 20 µg/mL (Fig. 4). After 4 hours of exposure, Hm21RSΔ*avsA* had greater sensitivity than Hm21RS and Hm21RSKΔ*avsA*Tn*7*::*avsA* for all four concentrations. At 0.5 µg/ml To, the wild-type strain continued to proliferate, while at the higher concentrations (5 and 10 µg/ml To) Hm21RS and Hm21RSKΔ*avsA*Tn*7*::*avsA* decreased but remained at higher concentrations than Hm21RSΔ*avsA*. These data reveal that AvsA contributes to resistance against To, especially at higher concentrations.

**Figure 4.**
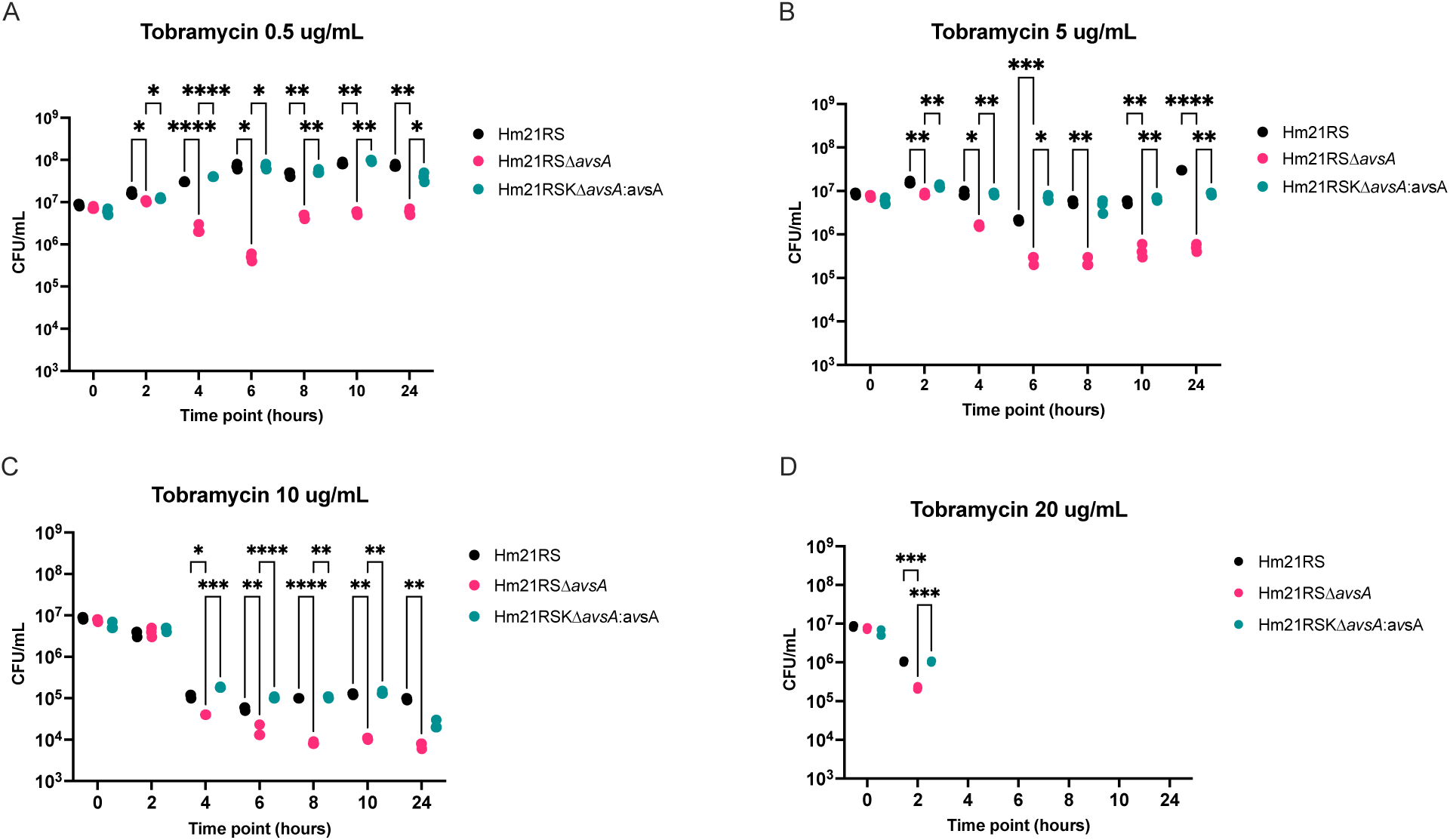
Loss of *avsA* Increases Tobramycin Susceptibility in *A. veronii* Hm21. Time-kill assay of *A. veronii* Hm21RS (black), ΔavsA (pink), and complemented Hm21RSKΔ*avsA*Tn*7::avsA* (green) strains exposed to increasing concentrations of with 0.5 µg/mL, 5 µg/mL, 10 µg/mL and 20 µg/mL of tobramycin. Each data point represents the average CFU/mL from a triplicate set. Hm21RSΔ*avsA* consistently showed greater killing, especially at higher drug concentrations. Statistical analysis was performed using one-way ANOVA, with *p < 0.05 and **p < 0.01.

Ciprofloxacin is a prophylactic agent for *A. veronii* infections [48]. For Cp we selected the concentrations of 0.024, 0.048, 0.096 and 0.192 µg/ml. The CFU/mL of all strains in all concentrations declined during the first 4 hours, with Hm21RSΔ*avsA* exhibiting heightened susceptibility at 6 hours, as evidenced by a lower CFU than in Hm21RS and Hm21RSKΔ*avsA*Tn*7*::*avsA* (Fig. 5A-D). These differences were more pronounced at concentrations of 0.096 and 0.192 µg/mL, as Hm21RSΔ*avsA* consistently demonstrated a greater reduction in CFU/mL (Fig. 5C and 5D). Thus, AvsA plays a role in resistance to antibiotics and the complement restores antibiotic resistance.

**Figure 5.**
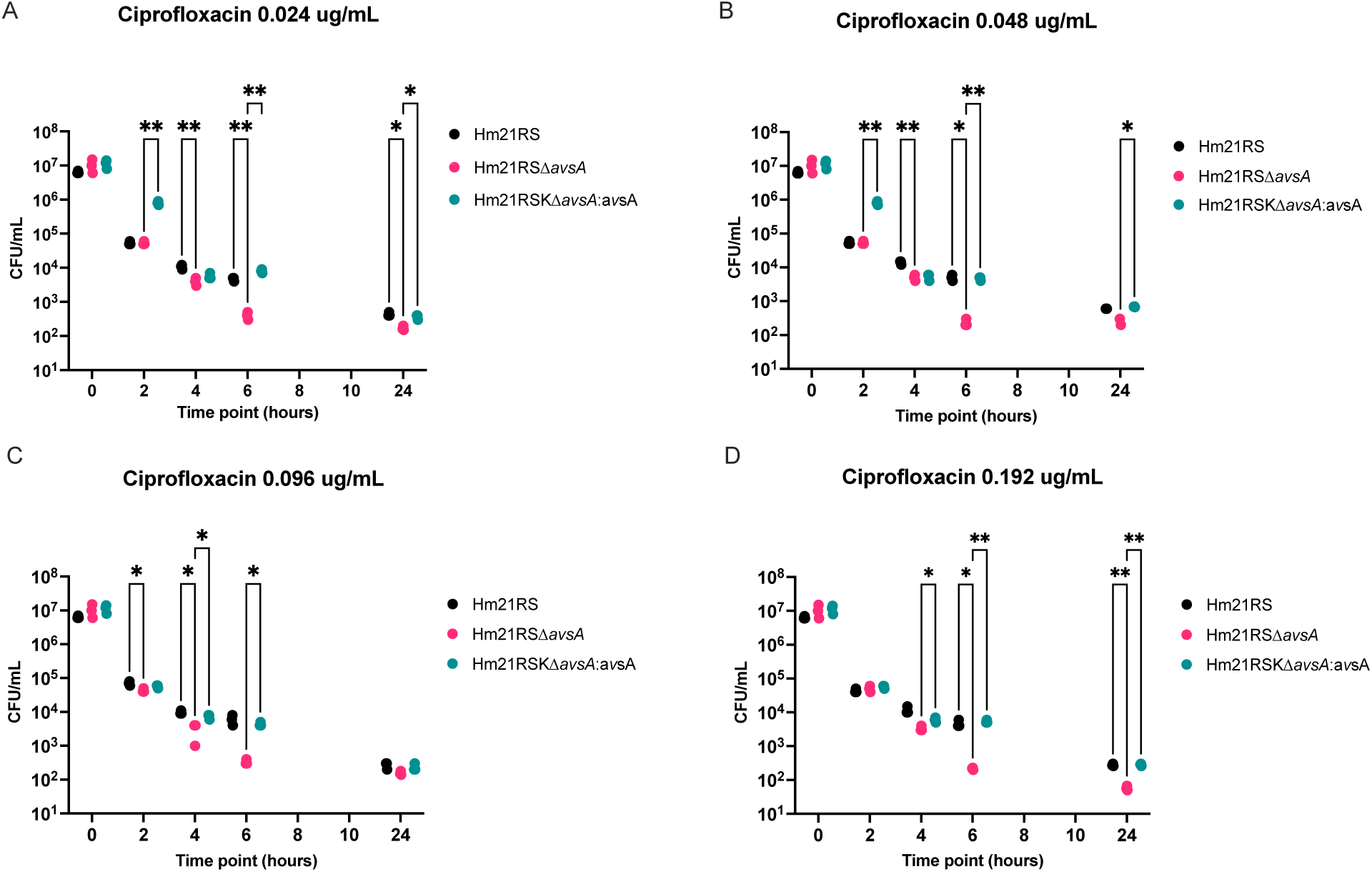
Deletion of *avsA* Enhances Ciprofloxacin-Mediated Killing in *A. veronii* Hm21RS. Time-kill assay of *A. veronii* Hm21RS (black), Hm21RSΔ*avsA* (pink), and complemented Hm21RSKΔ*avsA*Tn*7::avsA* (green) strains treated with ciprofloxacin at 0.024, 0.048, 0.096, and 0.192µg/mL over 24 hours. ΔavsA shows consistently lower survival, particularly at higher concentrations, while complementation restores resistance to wild-type levels. Points represent triplicate means; error bars show standard deviation. Statistical significance was determined by one-way ANOVA, with *p < 0.05 and **p < 0.01.

### AvsA contributes to colonization in the medicinal leeches

The ability of Hm21RSΔ*avsA* to colonize the leech digestive tract was assessed by competing this strain against a trimethoprim-resistant derivative of Hm21 and determining the competitive index in the intraluminal fluid (ILF) of the leech digestive tract 42h post-feeding [31], [49]. The median competition index for the Hm21RSΔ*avsA* in the ILF was around 0.4, which was significantly lower than that of Hm21RS and Hm21RSKΔ*avsA*Tn*7::avsA, Pvalues respectively here* (Fig. 6A). we used a blood control to assess whether the competitive disadvantage of the Δ*avsA* strain was digestive tract specific or is atributable to growth in heat-inactivated blood.. The same growth defect was observed in the blood control (Fig. 6B), in which the median competition index for the Hm21RSΔ*avsA* strain was around 0.1, which was significantly lower than that of Hm21RSKΔ*avsA*Tn*7::avsA* strains, *Pvalues respectively here*. This data indicates that the AvsA protein is important for fitness in the blood meal and thus also for growth inside the leech digestive tract.

**Figure 6.**
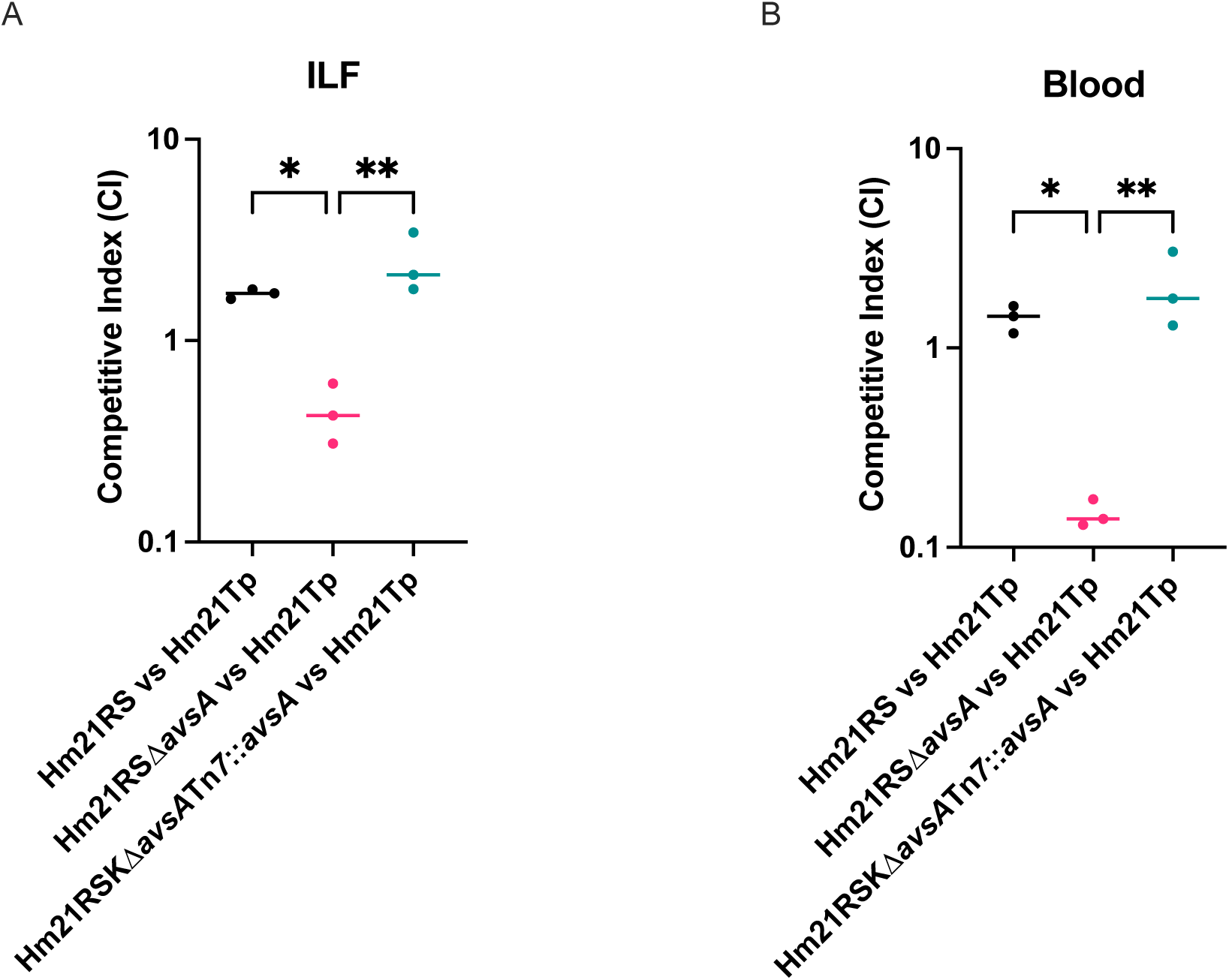
The AvsA protein is Essential for *in vivo* colonization Fitness of *A. veronii* Hm21RS. *In vivo* competition assays comparing Hm21RS (black), Hm21RSΔ*avsA* (pink), and Hm21RSKΔ*avsA*Tn*7::avsA* (green) strains in (A) ILF and (B) blood. Competitive index (CI) values below 1 indicate reduced colonization. Hm21RSΔ*avsA* shows a significant fitness defect in both environments, which is rescued by *avsA* complementation. Bars represent mean CI, and points show individual values. ANOVA and non-parametric t-tests were used for statistical analysis (*p < 0.05, **p < 0.01).

### The AvsA enhances biofilm formation in the Hm21 strain

The influence of surface layer proteins (SLPs) in biofilm formation has been well-documented, as these proteins play crucial roles in cell adhesion, environmental stability, and the structural integrity of biofilms [18], [50], [51]. To investigate the role of the SLP-encoding gene *avsA* in biofilm formation, biofilm-formation was measured in an M9 minimal medium with glucose (M9+Glu). Our results revealed that the Hm21RSΔ*avsA* exhibited a 50% reduction in biofilm (OD_590_ ± 0.0683) than the wild-type Hm21RS (OD_590_ ± 0.1027) and Hm21RSKΔ*avsA*Tn*7::avsA* (OD_590_ ± 0.1073), suggesting that the AvsA is important for biofilm formation (Fig. 7). To determine whether rifampicin and streptomycin resistance affect biofilm formation, the biofilm formation of the mother Hm21 strain was compared to rifampicin and streptomycin-resistant strains. The resistance to these antibiotics did not alter biofilm production (OD_590_ ± 0.1067). This indicates that the observed differences in biofilm formation between Hm21RS and Hm21RSΔ*avsA* are attributable to the *avsA* gene disruption rather than resistance mutations.

**Figure 7.**
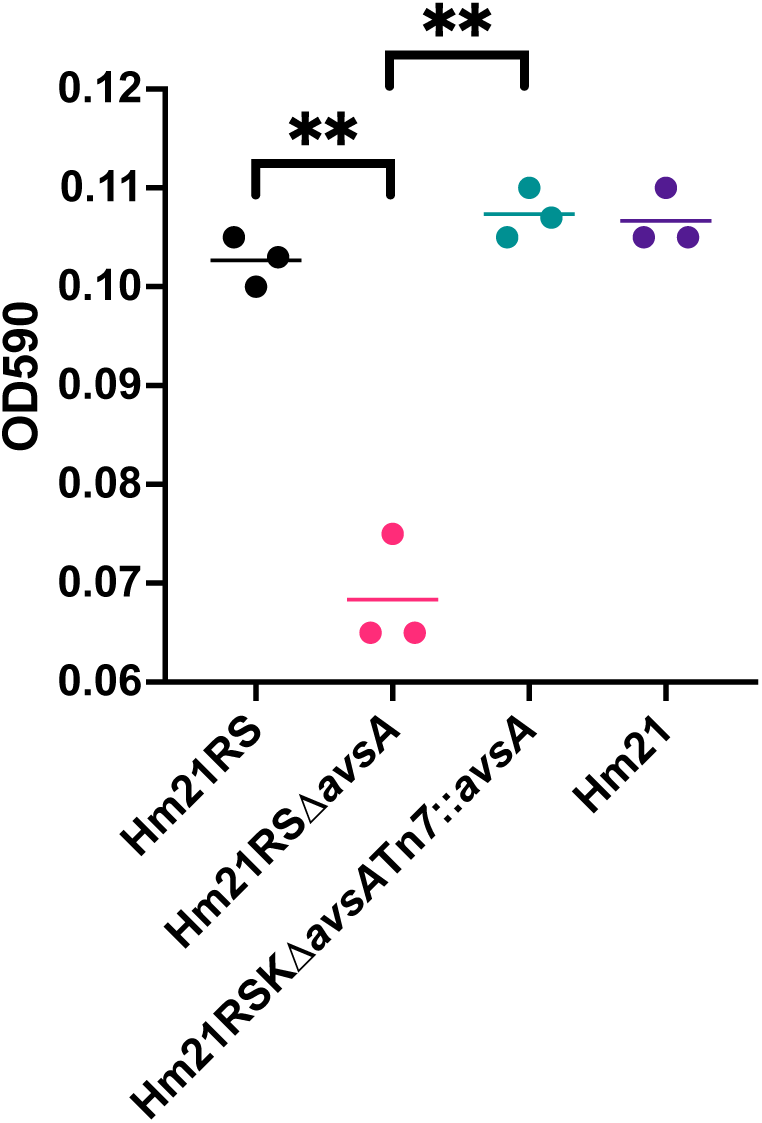
Biofilm formation abilities of Hm21RS, Hm21RSΔ*avsA*, Hm21RSKΔ*avsA*Tn*7::avsA*, and Hm21 strains. Biofilm formation was quantified by crystal violet staining and measuring OD₅₉₀ of adhered biomass. The Hm21RS*ΔavsA* exhibited significantly reduced biofilm formation compared to the Hm21*RS* strain. Complementation of the *avsA* (Hm21RSKΔ*avsA*Tn*7*::*avsA*) restored biofilm formation to wild-type levels. The parental strain Hm21 also showed similar biofilm levels as Hm21*RS*. Statistical analysis was performed using ANOVA to evaluate overall differences among the strains, followed by non-parametric t-tests to identify significant pairwise comparisons, with *p < 0.05 and **p < 0.01.

### Additional phenotypic tests

Sensitivities to sodium dodecyl sulfate (SDS) and sodium chloride were assessed in a minimal medium. The sensitivity to SDS was assessed in M9 medium containing 0.625%, 1.25%, 2.5% or 5% SDS. No difference in the growth curves of Hm21RSΔ*avsA* the was observed compared to the parent strain (Suppl Fig. 3). Similarly, no differences in the growth curves were observed in M9 medium containing 0%, 0.0625%, 0.125%, 0.25% and 0.5% NaCl (Suppl Fig 4), indicating that AvsA does not contribute to SDS or NaCl tolerance.

### Modeling of the *A. veronii* AsvA protein structure

We used AlphaFold 3 (AF3) [52] and EMSFold [53] to predict the structure of the protein. The AF3 model contains three distinct domains composed primarily of antiparallel beta strands with short, interspersed loop regions (Fig 8A and B). All three domains have antiparallel β-sheet β-sandwich motifs, with those of domains 1 and 2 reminiscent of the commonly found immunoglobulin-like fold. Overall, the pLDDT scores, which denote the confidence of the AF3 prediction, are high throughout the model (> 0.9) except for at the N- and C-termini of the protein, not unexpected particularly at the N-terminus where there is a secretion signal sequence. To corroborate this prediction, we also calculated a model using ESMFold, which takes into account evolutionary information based on multiple sequence alignments [53] (Fig. 8C). The domain folds from each program are similar, however the relative positioning of domains in each model was distinct. This variance may indicate flexibility between domains providing conformational variability to the protein.

**Figure 8.**
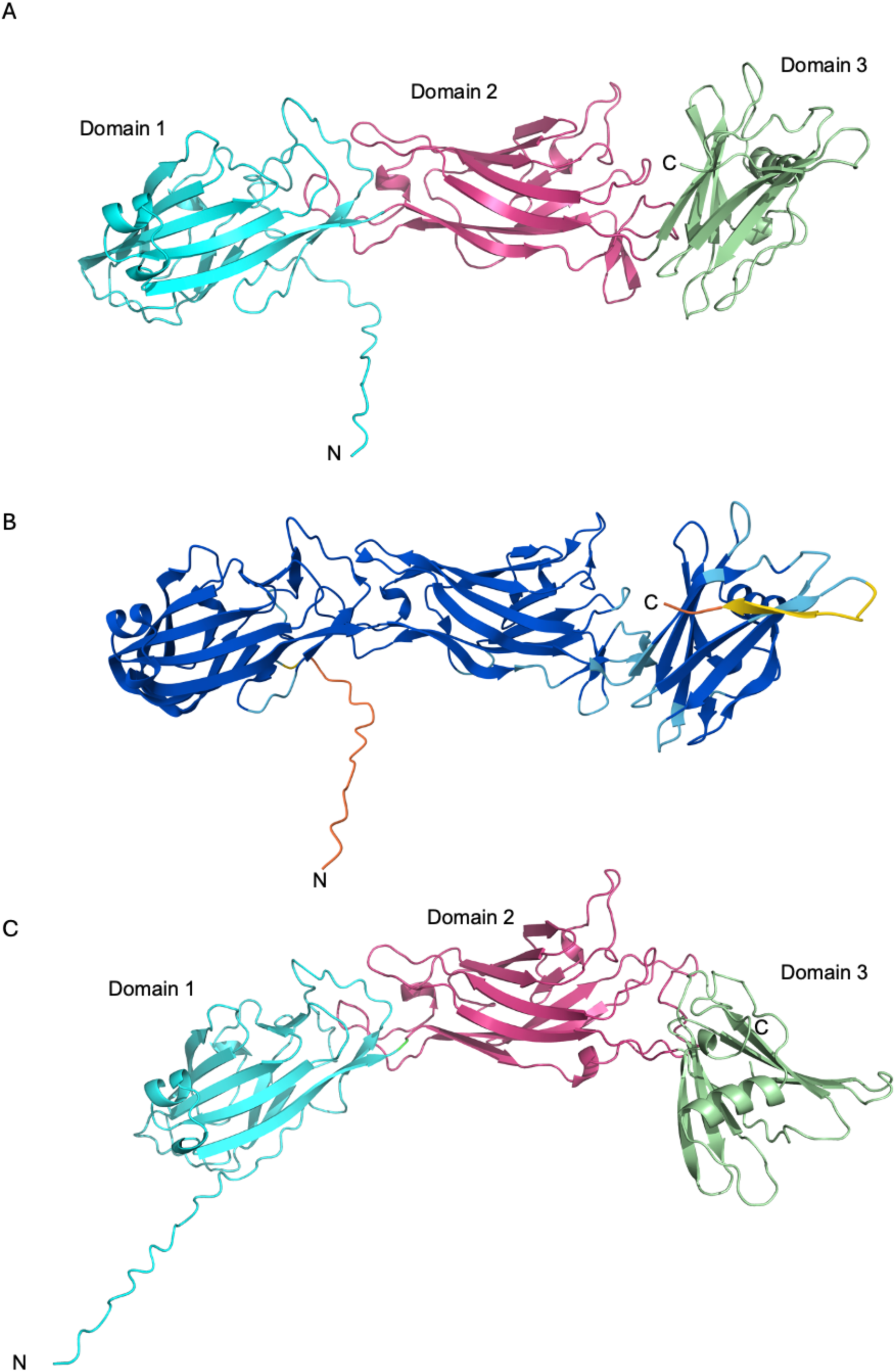
Ribbon representation of the structure of full-length *A. veronnii* AsvA as predicted with AlphaFold3 (AF3) and ESMFold. A) The AF3 model revealed three distinct domains, colored cyan, pink and green respectively. Domains 1 and 2 have immunoglobulin like folds (Ig(G)). B) The same AF3 model color-coded by the pLDDT values indicating the confidence level of the prediction. Most of the residues in the predicted structured have very high confidence scores (pLDDT > 90, colored dark blue), with the exception of the disordered secretion signal sequence at the N-terminus and a beta strand at the C-terminus (colored yellow) with pLDDT scores below 50. C) Ribbon diagram of the ESMFold prediction with similar structural features as the AF3 model. As in panel A, domains 1-3 are colored cyan, pink and green respectively. Visualization was done in PyMOL (The PyMOL Molecular Graphics System, Version 2.0 Schrödinger, LLC).

## Discussion

Our characterization of OMVs in *Aeromonas veronii strain* Hm21 resulted in the unexpected discovery of a widely distributed paracrystalline surface array protein, AsvA. AsvA covers the surface of both the bacteria and OMVs. SLPs on OMVs have only been described twice, in *Campylobacter fetus* [54], [55] and *Delftia acidovorans* [56]. In this study, we purified OMVs from *A. veronii* Hm21 and confirmed the presence of AvsA, a previously uncharacterized protein (M001_006550), by SDS-PAGE, TEM, and LC-MS (Fig. 1, Table 1). The SLP belongs to a homologous group found in over 300 strains, mostly described as hypothetical proteins, except for one from *A. hydrophila*. This lack of annotating these proteins properly, despite one well-characterized protein, is problematic as many researchers will miss important functional information. However, the detailed biochemical characterization of the SLP from *A. hydrophila* TF7 provides strong support that these proteins are SLPs (Fig. 2). This wide distribution of AsvA homologs suggests that the phenotypes we report could also be applicable to those organisms.

To determine the roles of the AvsA protein, we performed physiological assays, including growth curves, antibiotics, SDS and NaCl susceptibility, and biofilm formation, as well as an *in vivo* competition assay. Antibiotic sensitivity assays revealed a role of AsvA in resistance to tobramycin and ciprofloxacin. These findings are consistent with studies on *Clostridium difficile* SLPs protecting against antibiotics [46]. An *in vivo* competition assay showed Hm21RSΔ*avsA* had a colonization defect in the leech and a growth defect in blood, suggesting AvsA aids in overcoming barriers that are active in heat-inactivated blood and remain active inside the leech, perhaps antimicrobial peptides, heme toxicity or iron limitation. Most mutants that have been previously identified had a much more dramatic defect inside the leech digestive than in blood, e.g. *lpp, hgpB, hgpR,* and *exeM* mutants [31], [49], [57] but a mutation in a predicted GTP-binding protein had a similar phenotype [49]. Previous studies have shown that SLPs enhance biofilm formation by promoting bacterial adhesion to surfaces and strengthening cell-cell interactions, which leads to a stable biofilm matrix capable of withstanding environmental stresses [2]. Hm21RS*ΔavsA* mutant showed significantly reduced biofilm formation, suggesting that AvsA plays a crucial role in structural integrity and adhesion properties (Figure 7). Transmission electron microscopy (TEM) data suggest that AvsA is evenly distributed across the bacterial surface, indicating a uniform structural role rather than a localized function. This even distribution may support a hypothesis that AvsA contributes to surface protection in a manner analogous to other SLPs, such as those in *Bacillus anthracis* and *Clostridium difficile*, which have been implicated in resistance to host immune responses and antimicrobial agents [58].

The consistency of the observed phenotypes with other studies led us to assess if the predicted structure of AvsA is similar to other SLPs. Molecular models of AsvA revealed a three-domain structure composed of β-sandwich folds. In both models, the loops at the interface between domains 1 and 2 are intercalated suggesting they function as a singular, but malleable structural unit. The six-residue linker between domains 2 and 3 is extended resulting in a dissimilar orientation of these domains relative to one another in the models. Because SLPs are curved versions of 2D protein arrays, the conformational flexibility observed in the models may be important to assure correct assembly of the S-layer coat around the cell surface. Over the past decade, despite numerous issues associated with structural analyses of self-assembling molecular systems, there have been several structures of SLPs solved by both crystallography and cryo-EM and most recently cryo-ET. These include the S-layer in *Clostridioides difficile*, *Sulfolobus acidocaldarius, Lactobacillus acidophilus* and *Lactobacillus amylovorus, Haloferax volcanii, Geobacillus stearothermophilus* and *Caulobacter crescentus* [63][64][65][66][67][68]. Although they all share common core domains composed primarily of β-sandwich IgG-like domains, they vary greatly in size, amino acid composition and how they are modified post-translationally.

Therefore, how assembly is triggered, controlled and stabilized and how S-layers are anchored to the cell is still not understood. Structural studies of AsvA will provide great insight into these processes.

In summary, our study identified a widely distributed SLP that contributes to antibiotic resistance, host colonization, and biofilm formation. Notably, AvsA coats OMVs, raising important questions about its role in OMV function. For example, studies have demonstrated that OMVs contribute to biofilm development by facilitating the assembly of the biofilm matrix and promoting cell adhesion [65]. It will be important to determine whether AvsA enhances or alters these functions. Additionally, SLPs like AvsA may influence the ability of OMVs to fuse with other membranes. The interaction between OMVs and SLPs represents a compelling area for future investigation. Ongoing studies are focused on exploring how AvsA-associated OMVs may contribute to the modulation of colonization factors within a symbiotic host environment.

## Materials and Methods

### Bacterial Strains and Culture Conditions

The bacterial strains used in this study are derived from the rifampin and streptomycin-resistant *Aeromonas veronii* Hm21 (Hm21RS), as described in Rio 2007 [66]. All Hm21RS and Hm21RS-derived strains were cultured at 30℃ and *E. coli* strains were cultured at 37℃, both at 200 rpm with their respective antibiotics [67], [68], [69], [70], [71]. All strains were cultured in lysogeny broth (LB) (10 g/liter NaCl, 5 g/liter yeast extract, 10 g/liter tryptone) [72]. For phenotypic assays, Hm21RS and Hm21RS-derived strains were cultured in an M9 medium with 20% glucose (M9+Glu) [72]. Briefly, the M9 medium was prepared by first autoclaving the 878 mL of distilled water, followed by the addition of 100 mL of 10X M9 salts, 2 mL of 1M magnesium sulfate, 20 mL of 20% glucose, and 0.1 mL of 1M calcium chloride [72]. All growth media were supplemented with the appropriate antibiotics at the following concentrations: 20 ug/mL of Rifampicin (R), 100 ug/mL of Streptomycin (S), 100 µg/mL kanamycin (K), 100 µg/mL of trimethoprim (Tp), and plasmid maintenance with 100 µg/mL ampicillin.

### OMVs isolation and purification

To isolate outer membrane vesicles (OMVs) from *Aeromonas veronii* Hm21, we followed the protocol described by Seike group [43] with slight modifications. Cultures were centrifuged at 10,000 × g for 15 minutes at 4°C to pellet the cells. The supernatant was collected and filtered through a 0.22 µm pore-size filter (Millipore). The OMVs were concentrated by ultracentrifugation at 100,000 × g for 2 hours at 4°C. The OMV pellet was washed by resuspending it in phosphate-buffered saline (PBS) and centrifuging again under the same conditions. For further purification, the OMV pellet was resuspended in phosphate-buffered saline (PBS) and centrifuged in a discontinuous sucrose density gradient of layering 2 mL each of 10%, 20%, 30%, 40%, 50%, and 60% (w/v) and centrifugation at 100,000 × g for 17 hours at 4°C. The OMV band was collected from the 40-50% sucrose interface, diluted in PBS, and sucrose was removed with ultracentrifugation at 100,000 × g for 2 hours at 4°C. The purified OMVs were resuspended in PBS and stored at −80°C until further use.

### SDS-PAGE on OMVs

For the analysis of the OMV the SDS-PAGE protocol described by Seike et al. (2021) was followed with minor modifications. Purified OMVs were mixed with 4X Laemmli Sample Buffer (BIO-RAD, Hercules, CA) containing β-mercaptoethanol. Following denaturation at 95°C for 5 minutes the proteins were separated on a 12% SDS-PAGE with a constant voltage of 120 V. The gel was stained with Coomassie Brilliant Blue R-250 for 1 hour, followed by destaining with a solution of 10% acetic acid until clear protein bands were visible.

### Transmission electron microscopy preparation

Bacterial cultures were diluted 1:100 in fresh LB, collected, and centrifuged at 8,000 x g for 3 minutes to avoid breaking flagellar structures. The pellet was resuspended in 2.5% glutaraldehyde in 0.1 M cacodylate buffer (pH 7.2), and samples were fixed at 4°C for 2 hours. Sections were stained with 0.5% uranyl acetate in distilled water for 3 minutes, rinsed with distilled water, and air-dried. Stained sections were examined using FEI Tecnai 12 G2 Spirit Bio-Twin with AMT NanoSprint12 camera for transmission electron microscope (TEM) at the Bioscience Electron Microscopy Laboratory at the University of Connecticut at an accelerating voltage of 80-120 kV.

### Mass spectrometry analysis

OMVs were solubilized in 1 M urea and 0.1% RapiGest surfactant, followed by reduction of cysteine residues, alkylation, and complete enzymatic digestion with trypsin [73]. The resulting tryptic peptides were desalted and subjected to liquid chromatography-tandem mass spectrometry (LC-MS/MS) analysis on a Q Exactive HF mass spectrometer with a 1-hour gradient. Raw data files were searched against the complete Uniprot reference proteome for *Aeromonas veronii* Hm21 to account for potential bacteriophage capsid proteins. Identifications were filtered to achieve a 1% false discovery rate (FDR) at both peptide and protein levels using a target-decoy approach and reversed versions of the databases. FDR-filtered results were further analyzed using Scaffold 5 software (Proteome Software, version 5.3.3).

### Bioinformatic analysis of ORF M001_06550

The mass spectrometry data identified ORF M001_06550 from the *A. veronii* Hm21 genome as the most abundant protein in the OMV preparation. This ORF encoded a hypothetical protein and the deduced amino acid sequence was used as query for an NCBI blastp search of the NR database using default parameters, except that the number of allowable hits was increased to 250. The sequences were downloaded and further analyzed within the Geneious prime software package (version 2025.0.2). Sequences with an e-value ≤ 0.006 and that were longer than 300 amino acids were retained. The sequences were aligned using MAFFT and an approximate maximum-likelihood phylogeny with 100 local bootstraps was constructed using FastTree [74] and edited in iTOL [75]. The Signal sequences were identified using SignalP 5.0.

### Deletion and Complementation of *avsA*

Genomic DNA was extracted from overnight cultures using the QIAquick PCR Purification Kit (Qiagen), and the concentration and purity of the DNA were determined using Qubit Fluorometer (ThermoFisher) and Nanodrop (Thermo Fisher Scientific) spectrophotometers, respectively. To target the region flanking the *avsA* gene, two sets of primers were designed (Table 1), with tails to facilitate ligation via Gibson Assembly® (New England Biolabs) [76], [77]. The first primer pair (F1/R1_avsA_GA_pKAS46) amplified a 533 bp fragment upstream of the *avsA* gene, encompassing the promoter region and a tail for pKAS46. The second primer pair (F2/R2_avsA_GA_pKAS46) was designed to amplify a downstream region that included 57 bp of the *avsA* gene and extended to 438 bp downstreamand a tail for pKAS46. The DNA fragments were amplified using Phusion® High-Fidelity DNA Polymerase (New England Biolabs), and the resulting PCR products were purified using the QIAquick PCR purification kit (QIAGEN). The amplicons, equipped with internal tails for Gibson Assembly (New England Biolabs), were prepared for insertion into the multiple cloning site (MCS) of the suicide vector pKAS46 [70], which contains a kanamycin resistance marker.

The pKAS46 plasmid was linearized with *KpnI-HF* and *EcoRI-HF*, and purified by gel extraction and ethanol precipitation. The Gibson-assembled plasmid containing the a*vsA* deletion was transformed into chemically competent *E. coli* DH5α λ pir cells (New England Lab) .

Successful transformants were confirmed by Sanger sequencing (Plasmidasaur). Following conjugation on LB, merodiploid cells were selected by plating on LB Km_100_ agar. Subsequently, transconjugents were screened on LB for sensitivity to Sm to identify those colonies that lost the plasmid. To confirm the successful deletion of the gene, diagnostic PCR was performed using the Test_F/R_avsA primers. (Table 3).

**Table 2.** Strains and plasmids used in this study.

| Strain or plasmid | Description | Source |
| --- | --- | --- |
| ***A. veronii* Strains** |  |  |
| Hm21RS | Spontaneous Sm<sup>r</sup> mutant of Hm21R | [34] |
| Hm21Tp | Derivative of Hm21RS, Rif<sup>r</sup>, Sm<sup>r</sup> and Tp<sup>r</sup> | [57] |
| Hm21RSΔ*avsA* | Derivative of Hm21RS, *avsA* mutant | This study |
| Hm21RSKΔavsATn7:*avsA* | Hm21RS:ΔavsA containing Tn7Km<br>(ΔavsA:*avsA*Tn7Km), Rif<sup>r</sup>, | This study |
| ***E. coli* Strains** |  |  |
| WM3064-λpir | DAP auxotroph | [68] |
| CC118λpir | Carrying helper plasmid pEVS104 | [69] |
| Dh5α λpir | Chemically competent *E. coli* strain containing the *pir* genes | NEB |
| Dh5α::pKAS46:ΔavsA | Carrying the pKAS6 plasmid with the ΔavsA fragment | This study |
| WM3064 λpir::pUC18R6KT-mini-Tn7T:*avsA* | DAP auxotroph containing the pUC18R6KT-mini-Tn7T with the *avsA* gene and the promote region | This study |
| **Plasmids** |  |  |
| pKAS46 | pGP704, rpsL, Km<sup>r</sup> | [70] |
| pEVS104 | Helper Tra, Trb, Km<sup>r</sup>, Ap<sup>r</sup>, R6K replicon | [69] |
| pKAS46:ΔavsA | pGP704, rpsL, Km<sup>r</sup>, containing the ΔavsA fragment | This study |
| pUC18R6KT-mini-Tn7T | Ap<sup>r</sup>, R6K replicon, oriT, carrying Tn7 | [71] |
| pTNS3 | Helper plasmid encoding transposase gene for the chromosomal integration of mini-Tn7 | [67] |
| pUC18R6KT-mini-Tn7T:promoter+avsA | pUC18R6KT-mini-Tn7T carrying *avsA* gene and the promote region | This study |

**Table 3.**
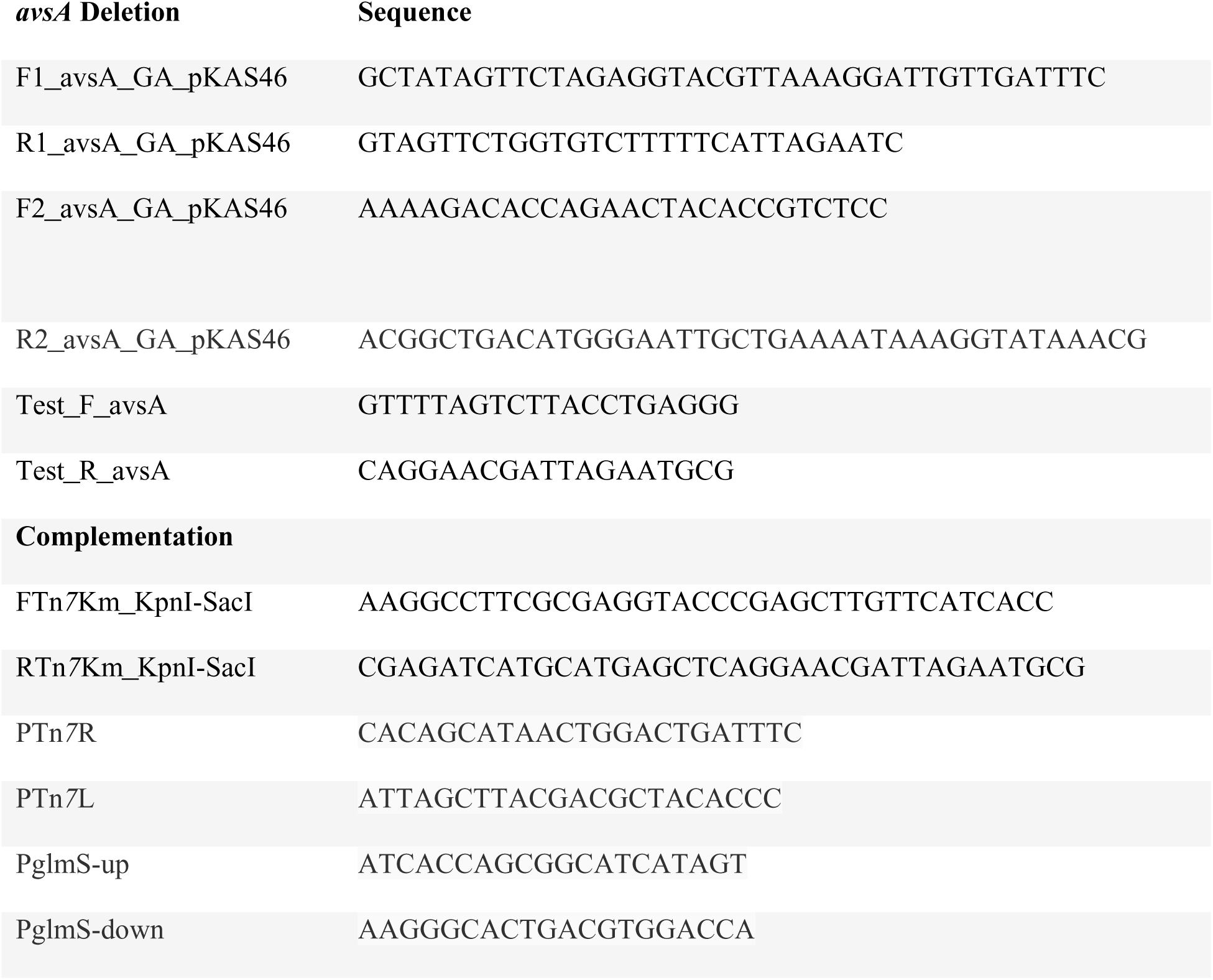
Primers used in this study.

Hm21Δ*avsA* was complemented using the pUC18R6KT-mini-Tn*7*T-Km vector [71]. The vector was double digested with *SacI*-HF and *KpnI*-HF. Primers designed for the amplicon included overhangs complementary to the linearized plasmid. Linear plasmid ligation was conducted using NEBuilder HiFi DNA Assembly, and the resulting plasmid was sequenced via Oxford Nanopore Technologies long-read Sanger sequencing (Plasmidasaur). This plasmid was then conjugated into chemically competent DH5α NEB 5-alpha E. coli (New England Biolabs) using a helper strain pEVS104 [69] and that contains the *pir* genes that are required for the R6K origin of replication. Subsequently, the plasmid was transformed into the E. coli strain WM3064 λ-*pir* [68], a diaminopimelic acid (DAP) auxotroph containing the *pir* genes essential for the R6K replicon. The resulting donor strain WM3064 λ-*pir*:pUC18R6KT-mini-Tn*7*T-Km-*avsA* was mated with the target strain Hm21RSΔ*avsA* using the *E. coli* helper pTNS3 that contains the transposase genes. Transformants were selected on LB agar containing 0.3 µM DAP. To select for Hm21RSΔ*avsA*Tn*7*::*avsA* strain, mating spots were streaked onto LB Km_100_ plates lacking 0.3 µM DAP. Complementation of the target strain was confirmed through colony PCR using primers described by Choi et al. [71], with modifications for the Hm21RS genome. These primers specifically target the Tn*7* and *glmS* regions, with confirmation further supported by Sanger sequencing.

### Antibiotic resistance testing

Overnight cultures grown in (M9+Glu), without antibiotics, were diluted to an optical density at 600 nm (OD_600_) of 0.01 in fresh M9 media. A series of antibiotic concentrations were prepared by diluting the antibiotics in fresh M9 media supplemented with glucose at twice the desired final concentration. In a 96-well plate, 100 µL of each bacterial strain was combined with 100 µL of the corresponding antibiotic concentration, resulting in the desired final antibiotic concentration in the wells. Optical density (OD_600_) measurements were taken every 2 hours for 52 hours using a spectrophotometer (BioTek).

To determine the minimum inhibitory concentration (MIC) of antibiotics for *Aeromonas veronii* Hm21, E-test strips (bioMérieux) were used following the manufacturer’s guidelines. E-test strips were placed on the surface of each agar plate. Plates were then incubated at 30°C for 18–24 hours. MIC values were recorded in µg/mL. Each test was conducted in triplicate.

### Antibiotic killing curves

The time-kill curves were performed as described by Foerster et al. and Montero et al. with modifications [78], [79]. 5 mL overnight cultures grown in minimal media M9+Glu were diluted to an OD_600_ of 0.3, corresponding to the mid-logarithmic growth phase, in 5 mL of fresh M9 supplemented with glucose. Tobramycin was added to the cultures to achieve final concentrations of 0.5, 5, 10, and 20 µg/mL, and ciprofloxacin at concentrations of 0.024, 0.048, 0.096 and 0.192 µg/mL. The colony-forming units per milliliter (CFU/mL) were determined at time zero for each strain to establish baseline bacterial counts. Cultures were incubated at 30℃ at 200 rpm, and aliquots were taken at 2, 4, 6, 8, 10 and 24 hours. These aliquots were diluted and spotted in triplicate onto LB agar plates, which were incubated for 18h at 30°C to determine bacterial viability at each time point. This assay was done in triplicate.

### *In vivo* Competition

A competition assay was performed with Hm21, Hm21RSΔ*avsA*, and Hm21RSKΔ*avsA:avsA* against Hm21Tp as described by Rio et al. and Silver et al. [49], [66]. The Hm21Tp strain, a Hm21 strain carrying trimethoprim (Tp) resistance, serves as the primary competitor [57]. Briefly, competing strains were plated and input CFU/mL was calculated on day one to confirm the input of 250 CFU/mL per strain. After a 42-hour incubation at room temperature (20°C), medicinal leeches *Hirudo verbana* (Leeches USA, LTD) were sacrificed, and the intraluminal fluid (ILF) was collected, diluted, and plated on selective media containing 100 ug/mL of trimethoprim at 100 ug/mL (Tp_100_) and Rif_20_ and Sm_100_ to quantify the output CFU/mL for each strain . The competitive index (CI) was calculated as follows:

CI = (mutant_output_/competitor_output_)/(mutantinput/competitor_input_).

### Biofilm Formation Assay

To quantify the biofilm formation of all Hm21 strains, we performed the protocol described by O’Toole [80]. The overnight culture was diluted to an OD_600_ of 0.01 at using a fresh M9 medium and 46-well plates were inoculated with 100 µL of the diluted cell suspension and 100 µL of fresh M9 medium per well. This was performed in triplicate. The plates were incubated at 30°C without shaking for 72 h, with daily media changes. After 72 h, biofilm structures were assessed by fixing the biofilm with ice-cold methanol for 15 minutes, followed by staining with 0.1% crystal violet solution (0.1 g of crystal violet in 100 mL distilled water) for 30 minutes. Excess dye was removed by gentle washing with distilled water, and the biofilm was then de-stained with 10% acetic acid for 30 minutes. The optical density of the de-stained solution was measured at 590 nm (OD_590_), with plain acetic acid used as a blank control.

### Growth curve using different concentrations of SDS

To evaluate the growth curve of Hm21RS and Hm21RSΔ*avsA* in the presence of detergents sodium dodecyl sulfate (SDS) and Triton X, we performed the protocol described by Silver et al. [49]. Overnight cultures were diluted to an OD_600_ of 0.01. 100 µL of the diluted culture was incubated with 100 µL of 5%, 2.5%, 1.25% and 0.625% of SDS in M9 in 96 well plates with a final cell concentration of 0.01 OD_600_ in each well. The plate was incubated at 30°C, and OD600 readings were taken every 2 hours by a spectrophotometer for 24 hours. This experiment was performed in triplicate.

### Osmotic assay using sodium chloride

Hm21RS and Hm21RSΔ*avsA* cells were cultured overnight in M9+Glu. Overnight cultures were diluted to a concentration of 0.01 OD_600_. 100 µL of the diluted cell cultures were incubated with 100 µL of NaCl for final concentrations of 0%, 0.0625%, 0.125%, 0.25% and 0.5% in 96 well plates. OD_600_ was measured every 2 hours for 18 hours.

### AlphaFold and ESMFold analysis of AvsA

A structure of full length AvsA was generated using AlphaFold3 [52]. The resulting coordinate file was compared with model prediction from ESMFold [53]. There was a general consensus about the folds of the three domains identified from both software packages however, the orientation of the domains relative to one another differed, particularly in regard to the positioning of domain 3. PyMOL was used to generate structural figures for the AlphaFold3 and ESM analyses (The PyMOL Molecular Graphics System, Version 2.0 Schrödinger, LLC).

### Statistical Analysis

All statistical analyses were performed using GraphPad Prism 9.0. For the analysis of *in vivo* colonization and biofilm formation, data were analyzed using one-way ANOVA followed by non-parametric *t*-tests to compare groups. Time-kill curve data were analyzed using two-way ANOVA, followed by non-parametric *t*-tests to assess differences across groups and time points. *p < 0.05 and **p < 0.01 were considered statistically significant.

## Acknowledgments

We want to express our deepest gratitude to Drs. Daniel Gage, Michele Maltz-Matyschsyk and Jeremiah Marden for their invaluable insights and expertise in phenotypic profiling and mutagenesis studies, which greatly contributed to the success of this research. We also extend our heartfelt thanks to Dr. Jeremy Balsbaugh for his assistance with the LC-MS/MS analyses provided by the Proteomics and Metabolomics facilities at the University of Connecticut (UConn). Additionally, we are grateful to the Bioscience Electron Microscopy Laboratory at UConn for their support and expertise in obtaining the TEM micrographs. Their collective contributions were instrumental to the completion of this work. This work was supported by U.S. Department of Agriculture (Award 8082-32000-007-000-D) and by the Chan Zuckerberg Initiative Science (Award 2022-253562-5022).

---

**Supplemental Fig. 1.**
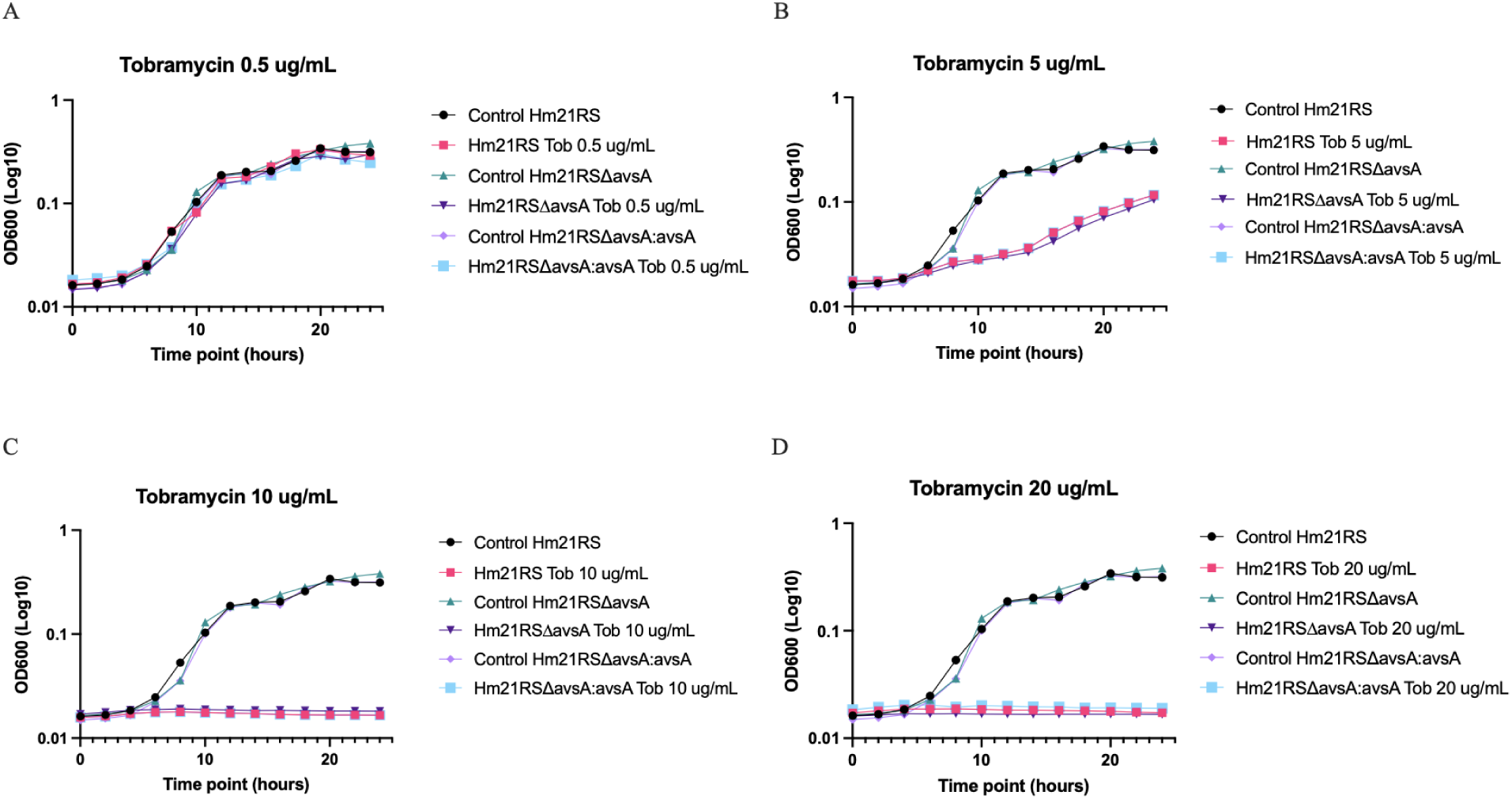
Growth curve in tobramycin of the Hm21RS and Hm21RSΔ*avsA* strains in M9 with 20% glucose.

**Supplemental Fig. 2.**
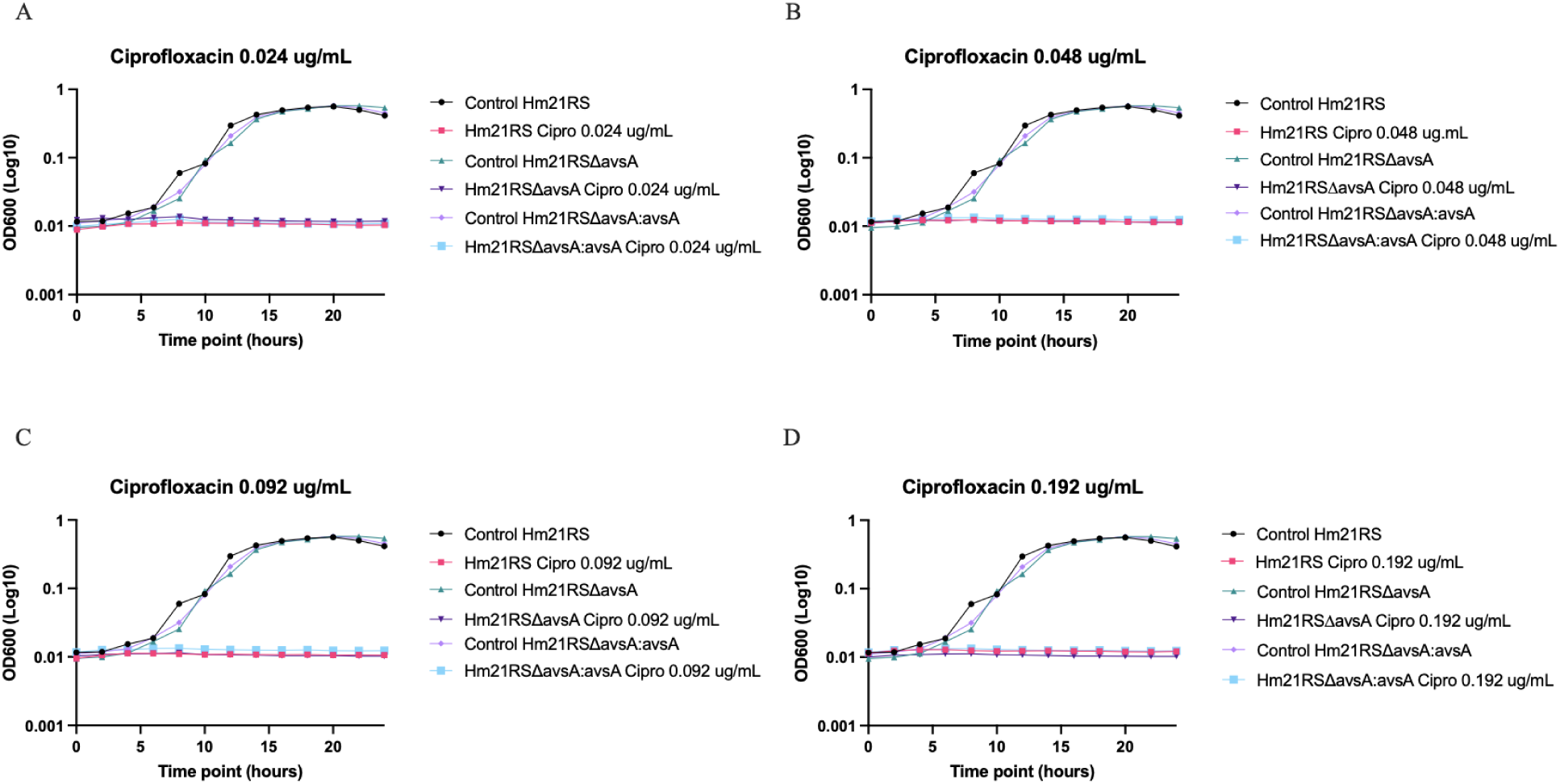
Ciprofloxacin sensitivity growth curve of the Hm21RS and Hm21RSΔ*avsA* strains in M9 with 20% glucose. Figure 3-A shows the overall growth curves treated with different concentrations ranging from 0.384 μg/mL to 0.003 μg/mL. Figure 3-B shows concentrations 0.096 μg/mL and 0.192 μg/mL, which suggests the Hm21RSΔ*avsA* is more sensitive than the Hm21RS strain.

**Supplemental Fig 3.**
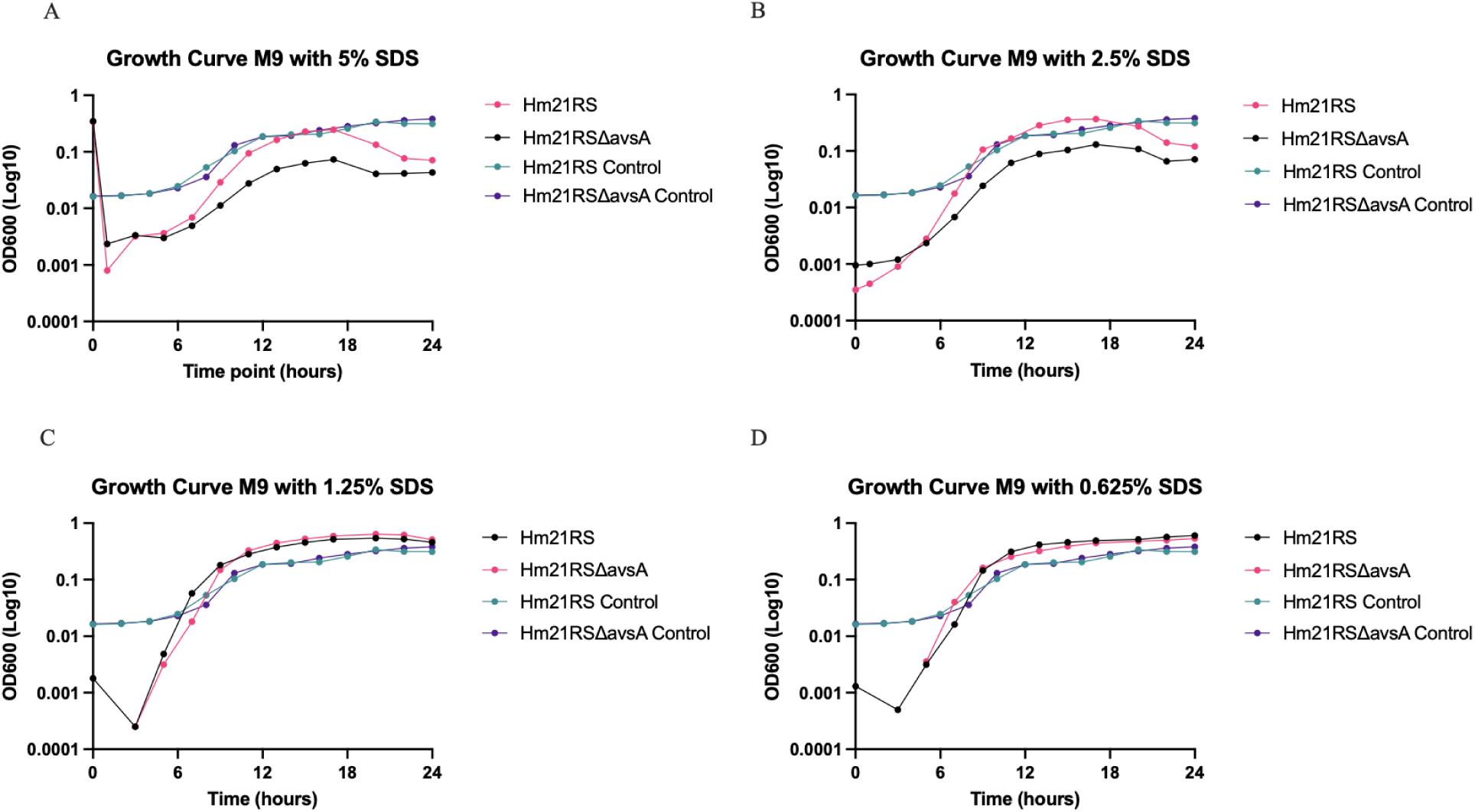
Effect of sodium dodecyl in the Hm21RS and Hm21RS*ΔavsA* strains. Growth curves of Hm21RS and Hm21RS*ΔavsA* strains in M9 minimal medium supplemented with varying concentrations of sodium dodecyl sulfate (SDS). Growth was monitored over 24 hours by measuring OD_600_ (log10 scale). Control conditions without SDS were included for each strain. (A) Growth in M9 with 5% SDS, (B) Growth in M9 with 2.5% SDS, (C) Growth in M9 with 1.25% SDS, and (D) Growth in M9 with 0.625% SDS. Hm21RS (pink), Hm21RS*ΔavsA* (black), Hm21RS *Control* (cyan), and Hm21RS*ΔavsA* Control (purple) are represented. Error bars indicate standard deviations of biological replicates.

**Supplemental Fig. 4.**
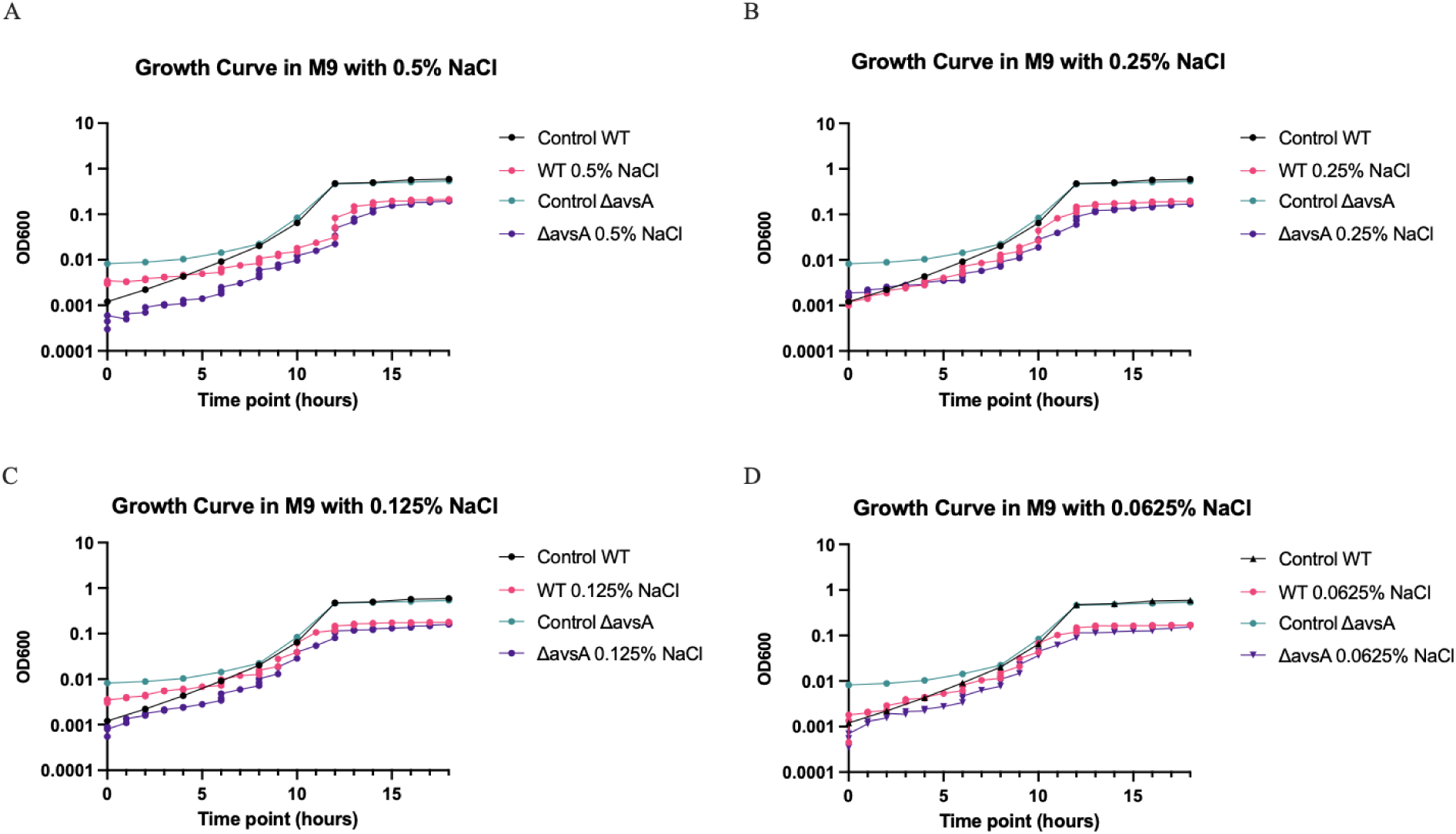
Growth curves of Hm21RS (WT) and Hm21*ΔavsA* (Δ*avsA*) strains in M9 minimal medium supplemented with varying concentrations of NaCl. Growth curves of Hm21RS (WT) and Hm21*ΔavsA* (*ΔavsA*) strains in M9 minimal medium supplemented with varying concentrations of NaCl. Bacterial growth was monitored over 18 hours. Control conditions without NaCl were included for each strain. (A) Growth in M9 with 0.5% NaCl, (B) Growth in M9 with 0.25% NaCl, (C) Growth in M9 with 0.125% NaCl, and (D) Growth in M9 with 0.0625% NaCl. WT *Control* (black), WT *+ NaCl* (pink), *ΔavsA* Control (cyan), and *ΔavsA +* NaCl (purple) are represented.

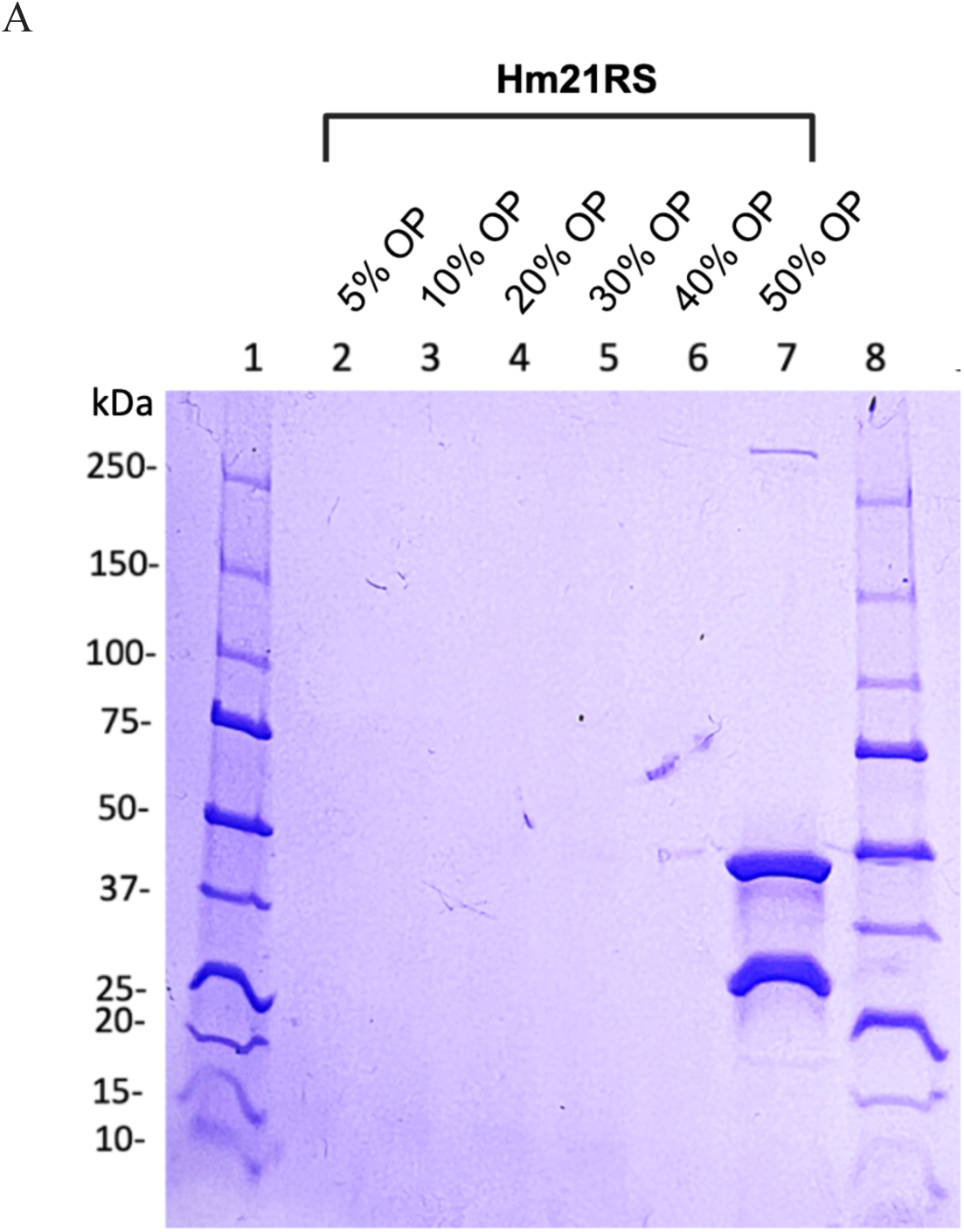

**Supplemental Table 1.** Total protein composition of outer membrane vesicles from *A. veronii* Hm21. <u>LC-MS Purified WT_avsA.xlsx</u>

**Supplemental Table 2.** Minimal inhibitory concentration (MIC) in µg/mL using E-strip.

| Antimicrobial Agent | Hm21RS | Hm21RSΔavsA |
| --- | --- | --- |
| Ciprofloxacin | 0.006 | 0.004 |
| Kanamycin | 4 | 4 |
| Polymyxin B | 2 | 2 |
| Polymyxin E | 3 | 4 |
| Tetracyclin | 0.064 | 0.064 |
| Gentamycin | 0.50 | 0.50 |

## References

[1] K. Murphy, “Influence of O Polysaccharides on Biofilm Development and Outer Membrane Vesicle Biogenesis in Pseudomonas Aeruginosa PAO1,” J. Bacteriol., vol. 196, no. 7, pp. 1306–17, 2014.

[2] T. J. Beveridge, “Functions of S-Layers,” FEMS Microbiol. Rev., vol. 20, no. 1–2, pp. 99– 149, 1997.

[3] A. V. Klieve, “Naturally Occurring DNA Transfer System Associated with Membrane Vesicles in Cellulolytic Ruminococcus Spp. of Ruminal Origin,” Appl. Environ. Microbiol., vol. 71, no. 8, pp. 4248–53, 2005.

4. J. W. Schertzer and M. Whiteley, “A Bilayer-Couple Model of Bacterial Outer Membrane Vesicle Biogenesis,” vol. 3, no. 2. p. 00297,11, 2012.

[5] S. Ayalew, “Proteomic Analysis and Immunogenicity of Mannheimia Haemolytica Vesicles,” Clin. Vaccine Immunol., vol. 20, no. 2, pp. 191–6, 2013.

[6] A. Olofsson, “Biochemical and Functional Characterization of Helicobacter Pylori Vesicles,” Mol. Microbiol., vol. 77, no. 6, pp. 1539–55, 2010.

[7] M. Yamaguchi, “A Porphyromonas Gingivalis Mutant Defective in a Putative Glycosyltransferase Exhibits Defective Biosynthesis of the Polysaccharide Portions of Lipopolysaccharide, Decreased Gingipain Activities, Strong Autoaggregation, and Increased Biofilm Formation,” Infect. Immun., vol. 78, no. 9, pp. 3801–12, 2010.

[8] S. Fulsundar, “Gene Transfer Potential of Outer Membrane Vesicles of Acinetobacter Baylyi and Effects of Stress on Vesiculation,” Appl. Environ. Microbiol., vol. 80, no. 11, pp. 3469–83, 2014.

[9] Y. Shen, “Outer Membrane Vesicles of a Human Commensal Mediate Immune Regulation and Disease Protection,” Cell Host Microbe, vol. 12, no. 4, pp. 509–20, 2012.

[10] S. W. Kim, “Outer Membrane Vesicles from Beta-Lactam-Resistant Escherichia Coli Enable the Survival of Beta-Lactam-Susceptible E. Coli in the Presence of Beta-Lactam Antibiotics,” Sci. Rep., vol. 8, no. 1, pp. 5402–0, 2018.

[11] C. Rumbo, “Horizontal Transfer of the OXA-24 Carbapenemase Gene Via Outer Membrane Vesicles: A New Mechanism of Dissemination of Carbapenem Resistance Genes in Acinetobacter Baumannii,” Antimicrob. Agents Chemother., vol. 55, no. 7, pp. 3084–90, 2011.

[12] H. M. Kulkarni, R. Nagaraj, and M. V. Jagannadham, “Protective Role of E. Coli Outer Membrane Vesicles Against Antibiotics,” Microbiol. Res., p. 181, 2015.

[13] H. I. Zgurskaya, “Trans-Envelope Multidrug Efflux Pumps of Gram-Negative Bacteria and their Synergism with the Outer Membrane Barrier,” Res. Microbiol., p. 169 7–8, 2018.

[14] U. B. Sleytr, “Crystalline Bacterial Cell Surface Layers (s Layers): From Supramolecular Cell Structure to Biomimetics and Nanotechnology,” Angew. Chem. Int. Edin Engl., vol. 38, no. 8, pp. 1034–54, 1999.

[15] B. M. Phipps, R. Huber, and W. Baumeister, “The Cell Envelope of the Hyperthermophilic Archaebacterium Pyrobaculum Organotrphum Consists of Two Regularly Arrayed Protein Layers: Three-Dimensional Structure of the Outer Layer,” Mol. Microbiol., vol. 5, no. 2, pp. 253–65, 1991.

[16] J. Mayr, “A Hyperthermostable Protease of the Subtilisin Family Bound to the Surface Layer of the Archaeon Staphylothermus Marinus,” Curr. Biol. CB, vol. 6, no. 6, pp. 739– 49, 1996.

[17] U. B. Sleytr and T. J. Beveridge, “Bacterial S-Layers,” Trends Microbiol., vol. 7, no. 6, pp. 253–60, 1999.

[18] M. Sara and U. B. Sleytr, “S-Layer Proteins,” J. Bacteriol., vol. 182, no. 4, pp. 859–68, 2000.

[19] J. Kern and O. Schneewind, “BslA, the S-Layer Adhesin of B. Anthracis, is a Virulence Factor for Anthrax Pathogenesis,” Mol. Microbiol., vol. 75, no. 2, pp. 324–32, 2010.

[20] A. Ryan, “A Role for TLR4 in Clostridium Difficile Infection and the Recognition of Surface Layer Proteins,” PLoS Pathog., vol. 7, no. 6, p. 1002076, 2011.

[21] K. Honma, “Role of a Tannerella Forsythia Exopolysaccharide Synthesis Operon in Biofilm Development,” Microb. Pathog., vol. 42, no. 4, pp. 156–66.

[22] S. F. Koval and S. H. Hynes, “Effect of Paracrystalline Protein Surface Layers on Predation by Bdellovibrio Bacteriovorus,” J. Bacteriol., vol. 173, no. 7, pp. 2244–9, 1991.

[23] S. Monette, “Massive Mortality of Common Carp (Cyprinus Carpio Carpio) in the St. Lawrence River in 2001: Diagnostic Investigation and Experimental Induction of Lymphocytic Encephalitis,” Vet. Pathol., vol. 43, no. 3, pp. 302–10, 2006.

[24] K. Krovacek, “Isolation, and Virulence Profiles, of Aeromonas Hydrophila Implicated in an Outbreak of Food Poisoning in Sweden,” Microbiol. Immunol., vol. 39, no. 9, pp. 655–61, 1995.

[25] S. Chu, “Structure of the Tetragonal Surface Virulence Array Protein and Gene of Aeromonas Salmonicida,” J. Biol. Chem., vol. 266, no. 23, pp. 15258–65, 1991.

[26] R. G. Murray, “Structure of an S Layer on a Pathogenic Strain of Aeromonas Hydrophila,” J. Bacteriol., vol. 170, no. 6, pp. 2625–30, 1988.

[27] R. Beaz-Hidalgo and M. J. Figueras, “Aeromonas Spp. Whole Genomes and Virulence Factors Implicated in Fish Disease,” J. Fish Dis., vol. 36, no. 4, pp. 371–88, 2013.

[28] J. M. Janda and S. L. Abbott, “The Genus Aeromonas: Taxonomy, Pathogenicity, and Infection,” Clin. Microbiol. Rev., vol. 23, no. 1, pp. 35–73, 2010.

[29] T. J. Wiles and K. Guillemin, “The Other Side of the Coin: What Beneficial Microbes can Teach Us about Pathogenic Potential,” J. Mol. Biol., vol. 431, no. 16, pp. 2946–56, 2019.

[30] M. C. Nelson and J. Graf, “Bacterial Symbioses of the Medicinal Leech Hirudo Verbana,” Gut Microbes, vol. 3, no. 4, pp. 322–31, 2012.

[31] M. Maltz and J. Graf, “The Type II Secretion System is Essential for Erythrocyte Lysis and Gut Colonization by the Leech Digestive Tract Symbiont Aeromonas Veronii,” Appl. Environ. Microbiol., vol. 77, no. 2, pp. 597–603, 2011.

[32] J. M. Bates, “Distinct Signals from the Microbiota Promote Different Aspects of Zebrafish Gut Differentiation,” Dev. Biol., vol. 297, no. 2, pp. 374–86, 2006.

[33] A. V. Banse, “Secreted Aeromonas GlcNAc Binding Protein GbpA Stimulates Epithelial Cell Proliferation in the Zebrafish Intestine,” Gut Microbes, vol. 15, no. 1, p. 2183686, 2023.

[34] J. Graf, “Symbiosis of Aeromonas Veronii Biovar Sobria and Hirudo Medicinalis, the Medicinal Leech: A Novel Model for Digestive Tract Associations,” Infect. Immun., vol. 67, no. 1, pp. 1–7, 1999.

[35] J. N. Marden, “Host Matters: Medicinal Leech Digestive-Tract Symbionts and their Pathogenic Potential,” Front. Microbiol., vol. 7, p. 1569, 2016.

[36] W. W. Kay, “Porphyrin Binding by the Surface Array Virulence Protein of Aeromonas Salmonicida,” J. Bacteriol., vol. 164, no. 3, pp. 1332–6, 1985.

[37] P. Doig, L. Emody, and T. J. Trust, “Binding of Laminin and Fibronectin by the Trypsin-Resistant Major Structural Domain of the Crystalline Virulence Surface Array Protein of Aeromonas Salmonicida,” J. Biol. Chem., vol. 267, no. 1, pp. 43–9, 1992.

[38] J. S. Dooley and T. J. Trust, “Surface Protein Composition of Aeromonas Hydrophila Strains Virulent for Fish: Identification of a Surface Array Protein,” J. Bacteriol., vol. 170, no. 2, pp. 499–506, 1988.

[39] B. M. Phipps and W. W. Kay, “Immunoglobulin Binding by the Regular Surface Array of Aeromonas Salmonicida,” J. Biol. Chem., vol. 263, no. 19, pp. 9298–303, 1988.

[40] T. K. Pham, “A Quantitative Proteomic Analysis of Biofilm Adaptation by the Periodontal Pathogen Tannerella Forsythia,” Proteomics, vol. 10, no. 17, pp. 3130–41, 2010.

[41] J. C. Caruana, S. N. Dean, and S. A. Walper, “Isolation and Characterization of Membrane Vesicles from Lactobacillus Species,” Bio-Protoc., vol. 11, no. 17, p. 4145, 2021.

[42] R. Prados-Rosales, “Isolation and Identification of Membrane Vesicle-Associated Proteins in Gram-Positive Bacteria and Mycobacteria,” MethodsX, vol. 1, pp. 124–9, 2014.

[43] S. Seike, “Outer Membrane Vesicles Released from Aeromonas Strains are Involved in the Biofilm Formation,” Front. Microbiol., vol. 11, p. 613650, 2021.

[44] E. D. Avila-Calderón, “The Outer Membrane Vesicles of Aeromonas hydrophila ATCC 7966: a Proteomics Analysis and Effet on Host Cells,” Front. Microbiol., vol. 9, no. 2765, 2018.

[45] S. R. Thomas and T. J. Trust, “Tyrosine Phosphorylation of the Tetragonal Paracrystalline Array of Aeromonas Hydrophila: Molecular Cloning and High-Level Expression of the S-Layer Protein Gene,” J. Mol. Biol., vol. 245, no. 5, pp. 568–81, 1995.

[46] Q. Zhou, “The cwp66 Gene Affects Cell Adhesion, Stress Tolerance, and Antibiotic Resistance in Clostridioides Difficile,” Microbiol. Spectr., vol. 10, no. 2, p. 0270421,21, 2022.

[47] Fuente-Nunez and Cesar, “The Bacterial Surface Layer Provides Protection Against Antimicrobial Peptides,” Appl. Environ. Microbiol., vol. 78, no. 15, pp. 5452–6, 2012.

[48] B. L. Jones and M. H. Wilcox, “Aeromonas Infections and their Treatment,” J. Antimicrob. Chemother., vol. 35, no. 4, pp. 453–61, 1995.

[49] A. C. Silver, “Identification of Aeromonas Veronii Genes Required for Colonization of the Medicinal Leech, Hirudo Verbana,” J. Bacteriol., vol. 189, no. 19, pp. 6763–72, 2007.

[50] L. L. Wong, “Surface-Layer Protein is a Public-Good Matrix Exopolymer for Microbial Community Organisation in Environmental Anammox Biofilms,” ISME J., vol. 17, no. 6, pp. 803–12, 2023.

[51] S. K. Shukla, T. Manobala, and T. S. Rao, “The Role of S-Layer Protein (SlpA) in Biofilm-Formation of Deinococcus Radiodurans,” J. Appl. Microbiol., vol. 133, no. 2, pp. 796–807, 2022.

[52] J. Abramson, “Accurate Structure Prediction of Biomolecular Interactions with AlphaFold 3,” Nature, vol. 630, no. 8016, pp. 493–500, 2024.

[53] Z. Lin, “Evolutionary-Scale Prediction of Atomic-Level Protein Structure with a Language Model,” in *Science*, New York, 2023, p. 379 6637.

[54] A. J. Winter, “Microcapsule of Campylobacter Fetus: Chemical and Physical Characterization,” Infect. Immun., vol. 22, no. 3, pp. 963–71, 1978.

[55] P. Farace, “Campylobacter Fetus Releases S-Layered and Immunoreactive Outer Membrane Vesicles,” Rev. Argent. Microbiol., vol. 54, no. 2, pp. 74–80, 2022.

[56] A. Shetty and W. J. Hickey, “Effects of Outer Membrane Vesicle Formation, Surface-Layer Production and Nanopod Development on the Metabolism of Phenanthrene by Delftia Acidovorans Cs1-4,” PloS One, vol. 9, no. 3, p. 92143, 2014.

[57] M. Maltz, B. L. LeVarge, and J. Graf, “Identification of Iron and Heme Utilization Genes in Aeromonas and their Role in the Colonization of the Leech Digestive Tract,” Front. Microbiol., vol. 6, p. 763, 2015.

[58] R. P. Fagan and N. F. Fairweather, “Biogenesis and Functions of Bacterial S-Layers,” Nat. Rev., vol. 12, no. 3, pp. 211–22, 2014.

[59] J. Herrmann et al., “A bacterial surface layer protein exploits multistep crystallization for rapid self-assembly,” Proc Natl Acad Sci U A, Jan. 2020, doi: 10.1073/pnas.1909798116.

[60] P. Lanzoni-Mangutchi, O. Banerji, and J. Wilson, “Structure and assembly of the S-layer in C. difficile,” Nat Commun, vol. 13, p. 970, 2022, doi: 10.1038/s41467-022-28196-w.

[61] L. Gambelli et al., “Structure of the two-component S-layer of the archaeon Sulfolobus acidocaldarius,” Elife, Jan. 2024, doi: 10.7554/eLife.84617.

[62] T. Sagmeister et al., “The molecular architecture of Lactobacillus S-layer: Assembly and attachment to teichoic acids,” Proc Natl Acad Sci U A, vol. 11;121(24):e2401686121, 2024, doi: 10.1073/pnas.2401686121.

[63] K. A, A. V, and B. TAM, “Complete atomic structure of a native archaeal cell surface,” Cell Rep, Nov. 2021, doi: 10.1016/j.celrep.2021.110052.

[64] E. Baranova, R. Fronzes, and A. Garcia-Pino, “SbsB structure and lattice reconstruction unveil Ca2+ triggered S-layer assembly,” Nature, vol. 487, pp. 119–122, 2012, doi: 10.1038/nature11155.

[65] W. Wang, W. Chanda, and M. Zhong, “The Relationship between Biofilm and Outer Membrane Vesicles: A Novel Therapy Overview,” FEMS Microbiol. Lett., vol. 362, no. 15, p. 117, 2015.

[66] R. V. M. Rio, M. Anderegg, and J. Graf, “Characterization of a Catalase Gene from Aeromonas Veronii, the Digestive-Tract Symbiont of the Medicinal Leech,” Microbiol. Read., vol. 153, no. Pt, pp. 1897–906, 2007.

[67] K.-H. Choi, “Genetic Tools for Select-Agent-Compliant Manipulation of Burkholderia Pseudomallei,” Appl. Environ. Microbiol., vol. 74, no. 4, pp. 1064–75, 2008.

[68] C. W. Saltikov and D. K. Newman, “Genetic Identification of a Respiratory Arsenate Reductase,” in Proceedings of the National Academy of Sciences of the United States of America 100.*19*, 2003, pp. 10983–8.

[69] E. V. Stabb and E. G. Ruby, “RP4-Based Plasmids for Conjugation between Escherichia Coli and Members of the Vibrionaceae,” Methods Enzymol., p. 358, 2002.

[70] K. Skorupski and R. K. Taylor, “Positive Selection Vectors for Allelic Exchange,” Gene, vol. 169, no. 1, pp. 47–52, 1996.

[71] K.-H. Choi, “A Tn7-Based Broad-Range Bacterial Cloning and Expression System,” Nat. Methods, vol. 2, no. 6, pp. 443–8, 2005.

[72] J. Sambrook and D. W. Russell, Molecular cloning: A laboratory manual. New York: Cold Spring Harbor, 2001.

[73] A. A. Klammer and M. J. MacCoss, “Effects of Modified Digestion Schemes on the Identification of Proteins from Complex Mixtures,” J. Proteome Res., vol. 5, no. 3, pp. 695–700, 2006.

[74] M. N. Price, P. S. Dehal, and A. P. Arkin, “FastTree: Computing Large Minimum Evolution Trees with Profiles Instead of a Distance Matrix,” Mol. Biol. Evol., vol. 26, no. 7, pp. 1641–50, 2009.

[75] I. Letunic and P. Bork, “Interactive Tree of Life (iTOL) V5: An Online Tool for Phylogenetic Tree Display and Annotation,” Nucleic Acids Res., vol. 49, no. W1, pp. 293– 6, 2021.

[76] D. G. Gibson, “Creation of a Bacterial Cell Controlled by a Chemically Synthesized Genome,” in *Science*, New York, N.Y, 2010, p. 329 5987.

[77] D. G. Gibson, “Enzymatic Assembly of DNA Molecules Up to several Hundred Kilobases,” Nat. Methods, vol. 6, no. 5, pp. 343–5, 2009.

[78] S. Foerster, “Time-Kill Curve Analysis and Pharmacodynamic Modelling for in Vitro Evaluation of Antimicrobials Against Neisseria Gonorrhoeae,” BMC Microbiol., vol. 16, pp. 216–9, 2016.

[79] M. M. Montero, “Time-Kill Evaluation of Antibiotic Combinations Containing Ceftazidime-Avibactam Against Extensively Drug-Resistant Pseudomonas Aeruginosa and their Potential Role Against Ceftazidime-Avibactam-Resistant Isolates,” Microbiol. Spectr., vol. 9, no. 1, p. 0058521,21, 2021.

[80] G. A. O’Toole, “Microtiter Dish Biofilm Formation Assay,” J. Vis. Exp. JoVE, vol. 47):2437, p. 10 3791 2437, 2011, doi: doi.47.

